# Working groups, gender and publication impact of Canada’s ecology and evolution faculty

**DOI:** 10.1101/2020.05.12.092247

**Authors:** Qian Wei, Francois Lachapelle, Sylvia Fuller, Catherine Corrigall-Brown, Diane S. Srivastava

## Abstract

A critical part of science is the extraction of general principles by synthesizing results from many different studies or disciplines. In the fields of ecology and evolution, a popular method to conduct synthesis science is in working groups – that is, research collaborations based around intensive week-long meetings. We present in this report an analysis of the impact of working group participation and gender on the publication impact of ecology and evolution faculty at Canadian universities who were research active over the last three decades (N=1408). Women are underrepresented in this research population relative to the general population, and even the Canadian faculty population. Participation in working groups not only benefits science, but also benefits the researchers involved by accelerating the temporal increase in their H-index. However, this benefit is particularly driven by senior male researchers. The effect is weaker for female researchers and even negative for researchers within 4 years of their PhD. However, gender does not affect current participation rates in working groups, nor reported indirect benefits – such as future collaborations, funding and data resources. The results of this study suggest that working groups can act as career catalysts for researchers, but that – as in many areas of science – there are challenging issues of equity that require action. Because the H-index is a cumulative measure, gender inequities from before the turn of the millenium may still be distorting the perceived publication impact of today’s research-active faculty.

## INTRODUCTION

Progress in science depends on our ability to draw out general principles from large amounts of heterogeneous data. While the grist of science is empirical observations and experimental manipulations - that is, primary research - the full strength of individual studies is only realized when they are synthesized, either statistically, mathematically or conceptually. Rapid development of computational tools to analyse large datasets, combined with the increasing availability of data through public repositories, has allowed scientists to harness the power of large-scale analyses of previously published results (Hampton et al. 2013). Such “synthesis science” allows researchers to determine which processes are truly general (by comparing multiple studies) and develop new paradigms (by exploring the interface between disciplines).

Much synthesis science is formally organized and funded through synthesis centers (Baron et al. 2017). These centres primarily fund working groups: a small network of researchers - typically 5 to 15 people - that meet to work intensively on a critical problem that requires the synthesis of large amounts of information, ideas or disciplines. Working groups are part of profound changes in how researchers conduct science. In many science disciplines, research is increasingly being conducted in collaborative networks – which can be seen in, for example, the impact of the Human Genome Project or The Intergovernmental Panel on Climate Change, or more generally in the rise in the number of authors per publication (Wuchty et al. 2007, Huang 2015). Such networks can achieve results that elude individual researchers (Baethge 2008). Research networks are able to apply more people-hours and a broader range of skills and expertise to collect or compile larger amounts of data and to do more complex or interdisciplinary analysis. Consequently, publications produced by research networks are higher impact than those produced by individual researchers (Wuchty et al. 2007). Research networks also often promote the types of knowledge exchange between countries that is valued by national governments (Hand 2010).

While it is apparent that scientific synthesis and working groups benefit science, the benefits to individual scientists are less clear. If Canada wants to support its researchers in achieving global impacts, we need to know how high impact research approaches affect scholars’ publication impact and careers - and whether this differs by gender and career stage. This information can then inform discussions of how the Canadian funding landscape might best support the careers of Canadian researchers to optimize impact and equity.

Women remain disadvantaged in multiple areas of science, for example in terms of grants (Urquhart-Cronish and Otto 2019, Witteman et al. 2019), citations (Huang et al. 2020) and hiring (Rivera 2017). Working groups and other collaborative networks that are more diverse in gender are more productive (Bear and Woolley 2011, Hampton and Parker 2011). However, this need not mean that there is gender equity in either the roles or recognition of the participants. For example, analyses of authorship contribution statements reveal that women are overrepresented in data collection roles and men in authorship roles (Macaluso et al. 2016). The “Matilda effect” (Rossiter 1993) refers to the systematic discounting of the quality or importance of contribution made by women scientists, as revealed in randomized experiments with scientific abstracts (Knobloch-Westerwick et al. 2013) or job applications (Moss-Racusin et al. 2012). Finally, women may participate less in working groups because of gendered childcare responsibilities – known generally to reduce women’s participation in international research collaborations (Uhly et al. 2017). All of the above may result in gender-specific impacts of working groups on careers.

## METHODS

We have constructed three databases to explore the impact of working groups on career advancement and gender equality in the EE faculty community in Canada: a Researcher Database, a Publication Database, and a Researcher Survey. This section will provide a detailed overview of these databases as well as the statistical methods we use for analyses.

### Researcher database

The study population in the Researcher Database includes all Canadian university faculty who received an NSERC Discovery grant through the Evolution and Ecology evaluation committee from 1991 to 2019 (N=1,408). We focus on this subset of scientists for two reasons: (1) biology as a discipline, including the ecology and evolution community, is more successful at integrating women than the physical sciences and engineering (although gender disparity remains). This ensures adequate sample size to conduct meaningful statistical analysis on gender differences; (2) focusing on one sub-discipline controls for disciplinary differences.

Our researcher database uses each individual researcher as the unit of analysis, providing a wide range of up-to-date (as of 2019) information. All information was manually collected from publicly available sources. These online sources, from the most preferred to the least preferred, were: Curriculum Vitae, websites of the researcher’s current institution, personally-maintained website of the researcher, websites of the researcher’s former institution(s), LinkedIn, Research Gate, Google Scholar, and other sources such as obituaries. Missing data are unavoidable since some information is not available from open sources.

We obtained information on the following fields, plus additional fields not used in this study (Appendix 1):

#### Gender

The gender of researchers was determined through analysis of first and middle names (if these names were strongly gender-associated), and inference of gender from photographs or pronouns on current websites. Such methods assume gender from external characteristics in the absence of direct information on researchers’ gender identity, and are therefore subject to some error. In particular, we may have misgendered researchers who identify as non-binary.

#### Academic degrees: Year of Completion and Granting institution(s)

We collected information on the year of completion and institution for Bachelor, Master and PhD degrees or their equivalents. In some cases, multiple web sources were needed to determine both the year and the granting institution. As the year of PhD completion was essential for our analysis, we were particularly thorough in validating this information, using searches of thesis databases maintained by academic institutions when necessary.

#### Current and previous institutions and departments

We recorded the name of the institution and department where the researcher is currently working, typically from institutional websites. To find previous institutions and departments, we relied on CVs and researcher-maintained websites.

#### Email addresses

*We* collected email addresses for as many researchers as possible to enable us to invite researchers in our focal population to complete our Researcher Survey.

#### H-index through time

We reconstructed the H-index of the researcher through time after building a Publication Database (described next).

#### Working group publications through time

We identified publications from the Publication Database that originated from working groups by automating the matching of titles and keywords followed by manual validation (described shortly).

### Publication database

The objective was to build a full retrospective publication record for all researchers for whom we successfully collected data from Web of Science Core Collection (hereafter, WOS). From the constructed publication record, the end-goal was to generate a longitudinal H-index in the Researcher database to measure the effect of synthesis science participation on the scientific impact of our scholars’ research output over time. To create a longitudinal H-index, we collected two groups/types of data: (1) retrospective publication record of peer-reviewed articles (as complete as possible) and (2) yearly distribution of citation counts. Although there are many freely available sources to collect individual scholars’ full publication record (CV, institutional/personal website, Google Scholar Profiles (GSP)), data availability is quite different when it comes to creating a longitudinal profile of a given scholar’s publication record. To our knowledge, there is no free, publicly available, and readily formatted data source that captures the H-index over time at the individual-researcher level. Therefore, we decided to use WOS for our data needs. As a curated collection of scientific scholarly content, WOS’s level of bibliometric coverage of scientific literature is very similar to that offered by other licensed indexing services (McLevey et al. 2018).

#### Collecting Web of Science bibliometric meta-data for raw publication list

There are several challenges in developing a clean list of publications from each scholar in our database. First, because WOS uses initials of all names but the surname, there is a likelihood of erroneously including publications by other scholars with similar names (e.g. Jane Doe not John Doe), a type of false positive. Second, researchers with middle names can vary between publications in which initials they include in their name (e.g. a WOS search on JM Doe would not include publications under J Doe), a type of false negative. Our workflow provides a means to reduce both the false positives and false negatives within the constraints of WOS. While we were completing our publication data collection in the summer of 2019, WOS introduced a beta search engine to better identify unique scholars but our work predates this.

We built an in-house tool in native Python computer language to automate the downloading of WOS publication and citation meta-data for each scholar for which we had a known PhD graduation year (n=1,247; 90%). The main advantages of programmatically carrying repetitive tasks is two-fold; time-efficiency and prevention of human imputation error. For each valid scholar, in the WOS Core Collection search engine, our Python program entered two pieces of information; (1) the scholar’s name - full last name, first initial of first name followed by the wild-card symbol (e.g., Zhang, Y*) and (2) year coverage - {PhD graduation year - 5} to 2019 (e.g., {1992-5} to 2019). Year coverage is called timespan by WOS. When using the wildcard symbol, WOS will return articles published by all the scholars whose last name is Zhang and first name starts with Y. Therefore, the resulting download for each query is not the exact publication/citation records of our very scholar, but rather a raw download of all the publication/citation records published by scholars whose names matched our queried name during the specified timespan. For each case, after the Python program has consolidated all the downloaded TXT files together, the resulting data format is a CSV file with publication as row-unit and relevant meta-data (author’s names, author’s institutional affiliation(s), year of publication, yearly citation counts, etc) as a separate column. In our case, by using the wildcard symbol and thereby expanding the range of results, the strategy was to ensure coverage of the potential variations in academics’ usage of initials in publication (e.g., John Wu; John S. Wu). Once completed, this raw download of WOS data yielded 1,269 CSV files totalling 640,289 raw publications (i.e. an average of 504 publications per researcher).

#### Filtering raw Web of Science bibliometric meta-data to generate a clean publication list

Cleaning raw WOS bibliometric data with the goal of generating a valid publication record for a specific scholar is a non-trivial task. We use a record linkage, rule-based algorithmic approach developed in Python to reduce/filter raw CSVs to clean CSVs. In a nutshell, our Python code uses information pertaining to (1) researchers’ name(s) as they appear in published works, (2) institutions where researchers were or are affiliated, and (3) publication titles collected outside the WOS platform to filter out as much false positive WOS publications as possible. Also called merge/purge processing or list washing (Hernández and Stolfo 1995), record linkage refers to a methodological strategy/process of joining records from one data source with another that describe the same entity. In this article, we refer to this process as a filtering process.

Since this filtering process relies on data collected outside the WOS corpus – i.e., publication history (CV, GSC, full list of publications, institutional trajectory) – we had to divide our 1,268 cases into four batches or categories based on the types and amount of data available for the record linkage/filtering code. The first category includes scholars for which we only successfully retrieved a CV that comprises a full list of publications (n=63; 5%; modal PhD year of 1991). The second category includes scholars for which we only found (and manually validated) an existing Google Scholar Profile populated with a list of publications (n=584; 46%; modal PhD year of 2000). The third category includes scholars for which we found both CV- and GSC-level data (n=167; 13%; modal PhD year of 2001). The four and final category includes scholars for whom we found no full or partial publication record outside WOS (n=452; 36%; modal PhD year of 1972). Since our sampled population covers several cohorts (PhD graduation year goes from 1953 to 2019), it is not surprising to find generational differences in online data availability.

Obviously, batch 4 scholars (n=452) represented an immense challenge in terms of the record linkage/cleaning process because of the lack of data-points on name spellings, publications, and academic history. In fact, for nearly half of batch 4 (n=197; 44%) we had to generate the clean WOS publication record relying on a mix of automation and manual work; namely, running the Python program, manual inspection of the CSV output, and repeated online searches for additional data-points. Our cleaning process is described in detail in Appendix 2, but in brief involves four filtering steps. The first filtering step matches known variants in the researcher’s name in publications (from online CV or Google Scholar profile) to those in the author fields in the raw WOS data. The second filtering step matches known institutional affiliations (past and current) from our Researcher Database to those in the raw WOS data. The third filtering step uses fuzzy matching of publication titles from the online CV and/or GSC profile with the title field of the raw WOS data. The fourth step is a dynamic, recursive process of inferring other affiliations for each researcher that were hitherto unknown to us, either by collecting this information when researchers have multiple affiliations listed on a WOS publication, or by harvesting the information from the GSC profiles. These additional institutions can then expand the second step of filtering by institution.

#### Computing H-Index

After completing the secondary cleaning of the filtering process, we generated the retrospective h-index using the open-source Python package “hindex” for the final corpus of WOS publications. The only manipulation we must operate on WOS yearly citation count is to transform the count from a yearly non-cumulative to a yearly cumulative one.

#### Identifying researcher publications originating from working groups

Once the publication corpus and its associated longitudinal h-index portrait is generated, the final task in extending the researcher database is to identify which publications are products of working groups. To achieve this level of granularity in the publication record, we devised approaches to capture publications originating from working groups funded by synthesis centers. We achieved this by matching WOS titles with known working group publications funded by synthesis centers (information from synthesis centre websites or obtained directly from centres), or by searching the funding and acknowledgement sections for synthesis centre names (in full or part) or their acronyms. We based our list of 15 synthesis centres on that published by the International Synthesis Consortium (https://synthesis-consortium.org/ accessed July 2019) supplemented by expert knowledge from a synthesis centre Director. We additionally captured working groups funded through other organizations or mechanisms, by searching for keywords commonly used to describe working groups (“working group”, “synthesis group”, “synthesis working group”, “synthesis committee”, “synthesis workshop”, “catalysis group”). All publications identified by the above methods were manually validated by two researchers experienced in biology and synthesis research, often by examination of the original article and perusal of synthesis centre websites. Full details of this process are given in Appendix 3.

#### Identifying publications as primary research, synthesis science and type of synthesis

For all of the publications identified as originating from a working group, two of our teammates with synthesis- and biology-related knowledge manually scored each title using meta-data from WOS (authors, abstract, funding, acknowledgements, abstract and WOS keywords) supplemented by inspection of original publications. Specifically, we scored each publication in terms of primary research vs. synthesis research, and as working group method vs. non-working group method.

Our team further categorized the synthesis research publications into the following types of synthesis work: statistical synthesis (involving the statistical analysis of previously published or archived data collected by multiple different researchers and/or studies), conceptual synthesis (qualitative review of the literature or proposal of new frameworks for scientific concepts or investigation), or mathematical synthesis (theoretical mathematical models or specific application of general models for the purpose of prediction).

For comparison, we scored the non-working group publications using similar criteria to the working group publications. We changed methods however, to allow for programmatic approaches to identify publications based on keywords given the large number of publications involved. We searched for keywords indicative of the three types of synthesis science (Appendix 4). This process yields 2,541 new WOS titles that had to be manually validated, 369 of which were actually primary research. We present this manually validated set in our report, rather than the entire set of non-working group publications as we are still actively developing machine learning methods to improve our automated scoring.

#### Creation of Publication Database

We created the publication database (n=80,777) by populating each row of a CSV file with all the unique publications contained in each researcher’s individual WOS publication record. We found a total of 17 260 publication duplicates (17.6%). In our context, a publication duplicate indicates that at least two of our sampled ecology and evolution researchers co-published a scientific article together. Given that our research studies a population of scholars working in the same field of expertise, the same country, and sometimes even the same department within a university, a level of redundancy is to be expected. Architecturally speaking, a publication-centric database is the native format of WOS publication and citation metadata. WOS data already comes formatted as such. Therefore, when producing our publication database we essentially merged all the individual publication record CVSs previously created for each scholar into one document while keeping every WOS publication metadata (i.e., year of publication, abstract, abstract keywords, yearly citation count, etc.). That said, we did conduct a number of transformations to the original WOS publication metadata in preparation for the analyses. For instance, we converted the yearly citation count [i.e. 1998:1, 1999:4; 2000:5] into a normalized format [year_1_since_pub: 1, year_2_since_pub: 4, year_3_since_pub: 5]. We also added a number of dummy variables in order to be able to carry our planned sets of statistical analyses. Since we were primarily interested in looking at the scientific impact differences between (1) types of research [primary vs. synthesis] and between (1) types of synthesis research [meta-analysis, systematic review, or mathematical modelling] respectively, we created five dummy variables: primary research, synthesis research, meta-analysis, systematic review, and mathematical model.

### Survey

An online survey (Appendix 5) was conducted to collect research data from EE faculty in Canada. We used our researcher database as the sample frame to recruit participants. As discussed above, the researcher database includes 1,408 EE faculty members in Canadian universities, and we were able to find 1,151 researchers’ email addresses. An email invitation was sent to all 1,151 scholars containing a link to an online questionnaire with a consent cover letter. If the questionnaire was completed, it was assumed that consent was given. We also collected each participant’s identification information for cross-referencing to researchers’ H-index in the extended researcher database.

Prior to distribution, the original questionnaire was pretested on three EE faculty members – one assistant professor, one associate professor and one full professor – and revised based on their feedback. The survey was carried out from July to September 2019. Two rounds of reminders were sent to improve the response rate. Eventually, 182 replies were received, amounting to a response rate of 15.8%. After clearing invalid questionnaires with too much missing data or no identification information, we have 169 valid responses, for an effective questionnaire response rate of 14.7%. In addition to questions such as academic progression and ethnicity, our survey data also provide nuanced information including whether researchers have declined working groups and why, their childbearing and parental leave.

### Statistical methods

We estimate regression models with fixed effects for individuals and years to assess the relationship between working groups and each researcher’s H-index over time and whether this effect differs for male and female researchers. Particular individuals may have unobserved characteristics (e.g. self-efficacy, social network) that lead both to high productivity and/or impact and that increase their chance of joining working groups. Differences in time periods may also matter for both patterns of research productivity and citation and working group participation. The fixed effects account for all unobserved time-invariant heterogeneity within individuals and common characteristics to all individuals in a specific year. In effect,each individual is treated as their own control (Allison 2009). In our case this means comparing researchers’ H-indices in years before and after participating in working groups, and then averaging those differences across the researchers we study. A Hausman test suggests that the fixed effects approach is favored over random effects regression. For any given individual, an H-index in one year is likely correlated with their H-index in previous and future years. In addition, there may be heteroscedasticity across individuals. To deal with these issues, we use the technique of clustering on individuals suggested by Wooldridge (2016) to obtain robust standard errors. We estimate our models based on subsets of the following specification of the full model: 

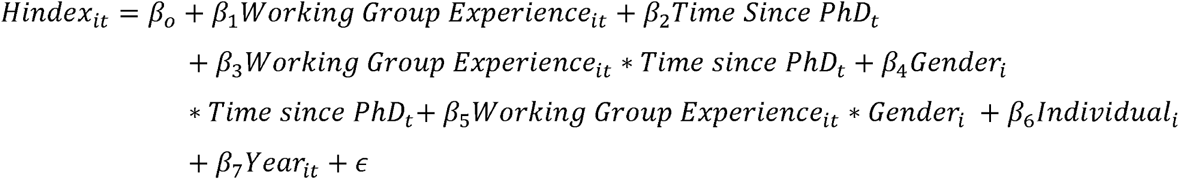

where *i* and *t* are index for individuals and years since PhD respectively; □ is the error term; *Individual*_*i*_ refer to the person-specific intercepts and *Year*_*it*_ are calendar year-specific dummy variables.

### Measures

#### Dependent variable

The key dependent variable in this study is researchers’ publication impact, which is measured by H-index. H-index is widely used in decision-making for academic hiring, university advancement, research awards and funding decisions. An H-index of *n* simply means that the researcher has published *n* papers that have been cited *n* times. The H-index is a cumulative measure, so can only increase with time, as shown for each of the 1247 faculty in our longitudinal database.

#### Independent variables

##### Time since PhD

The first independent variable is the time since each individual researcher received their PhD degrees. In our sample of the researcher population, built in 2019, this variable ranges from 0 to 69, comprising researchers who received their PhD from 1950 to 2019. We also transform this variable to account for the nonlinear relationship between time since PhD and researchers’ H-indices. Although these uncorrected trajectories appear almost linear, this is because overall citation rates have increased in recent decades. If we factor out the effects of calendar year, the H-index trajectories actually increase more slowly over time. A log-log model suggests the power is 0.6, so we transform this variable into (Time since PhD)^0.6^ to correct the power function relationship.

##### Working Group experience

This variable is coded as a dummy variable. If a researcher participates in a working group, they are coded as 1 for the year of the first publication from a working group and remain 1 after that experience; otherwise they are coded 0. In our sample, 85%(1,063 out of 1,247) of researchers have never participated in working groups and 15%have working group experience.

##### Gender

This variable is a dummy where female = 1 and male = 0. The percentage of female researchers in our research database is 23%.

## RESULTS AND DISCUSSION

### Ecology and Evolution Research Faculty in Canada

To understand the ecology and evolution (EE) research faculty population at Canadian universities, we utilize multiple sources of information – an online survey we conducted and a database of researchers and their publications that we built, as well as data drawn from Statistics Canada. We defined our study population as all researchers who held at least one NSERC Discovery Grant in the Evolution and Ecology committee between 1991and 2018 (N=1,408). We realize that some researchers, who otherwise consider themselves ecologists and evolutionary biologists, hold Discovery Grants at other committees, and that it is also possible to have a successful research career while never holding a Discovery Grant. Nonetheless, this is a simple and unambiguous criteria for defining the population of study.

Gender inequality still exists in the university faculty population in Canada. Overall, female researchers are underrepresented among university faculty (40.6%of all faculty): Figure 1). Women are even more underrepresented among EE faculty, as revealed both in our survey (33.1%) and researcher database (22.8%) (Figure 2). The difference in the proportion of female researchers between the survey and database reflects differences in the population sampled by each method: the survey was only completed by researchers with functional and public email addresses, generally younger (and therefore more likely to be female) researchers.

**Figure 1.**
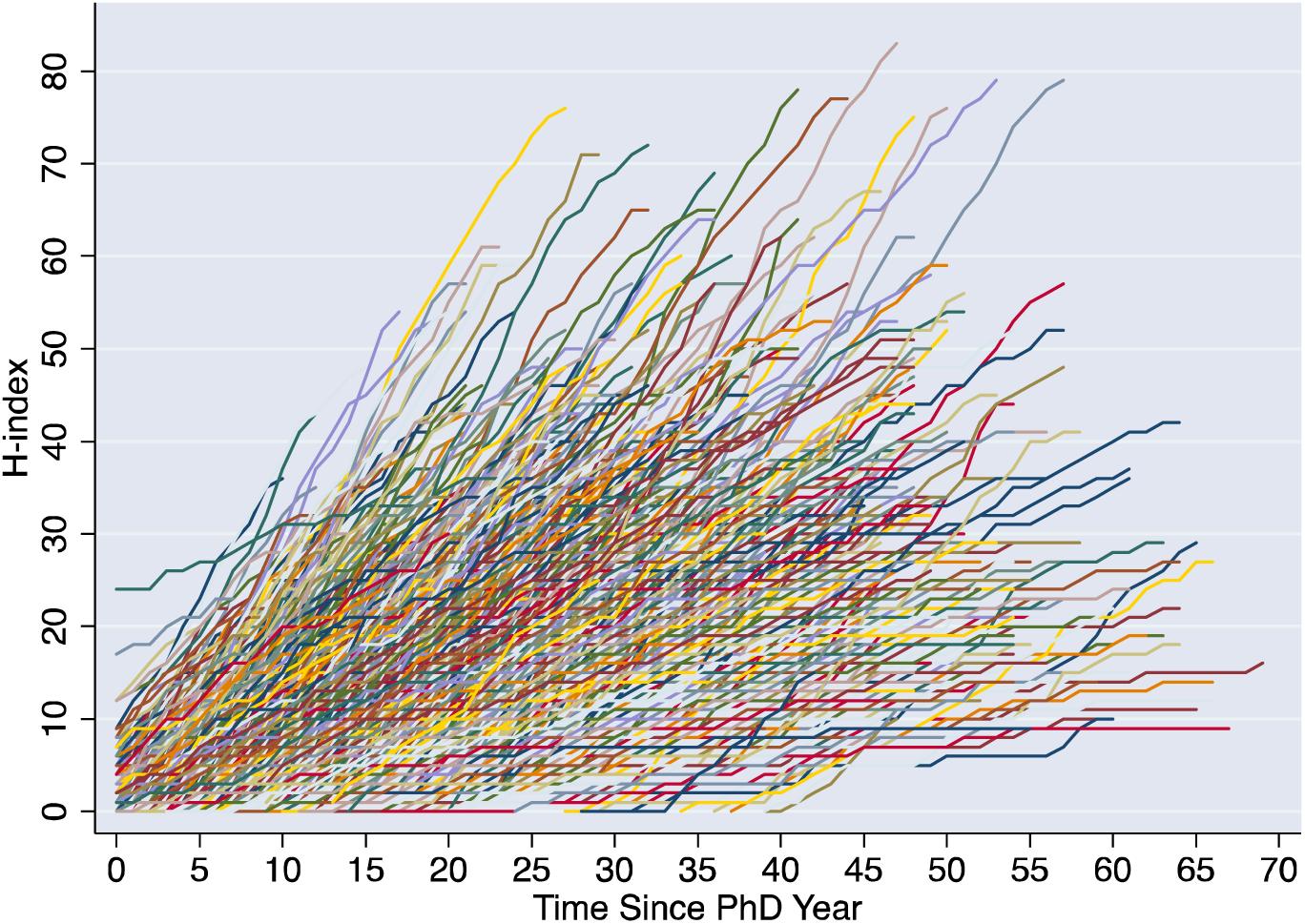
The H-index of each individual researcher (different coloured lines) increases with years since PhD. Source: Researcher Database

**Figure 2.**
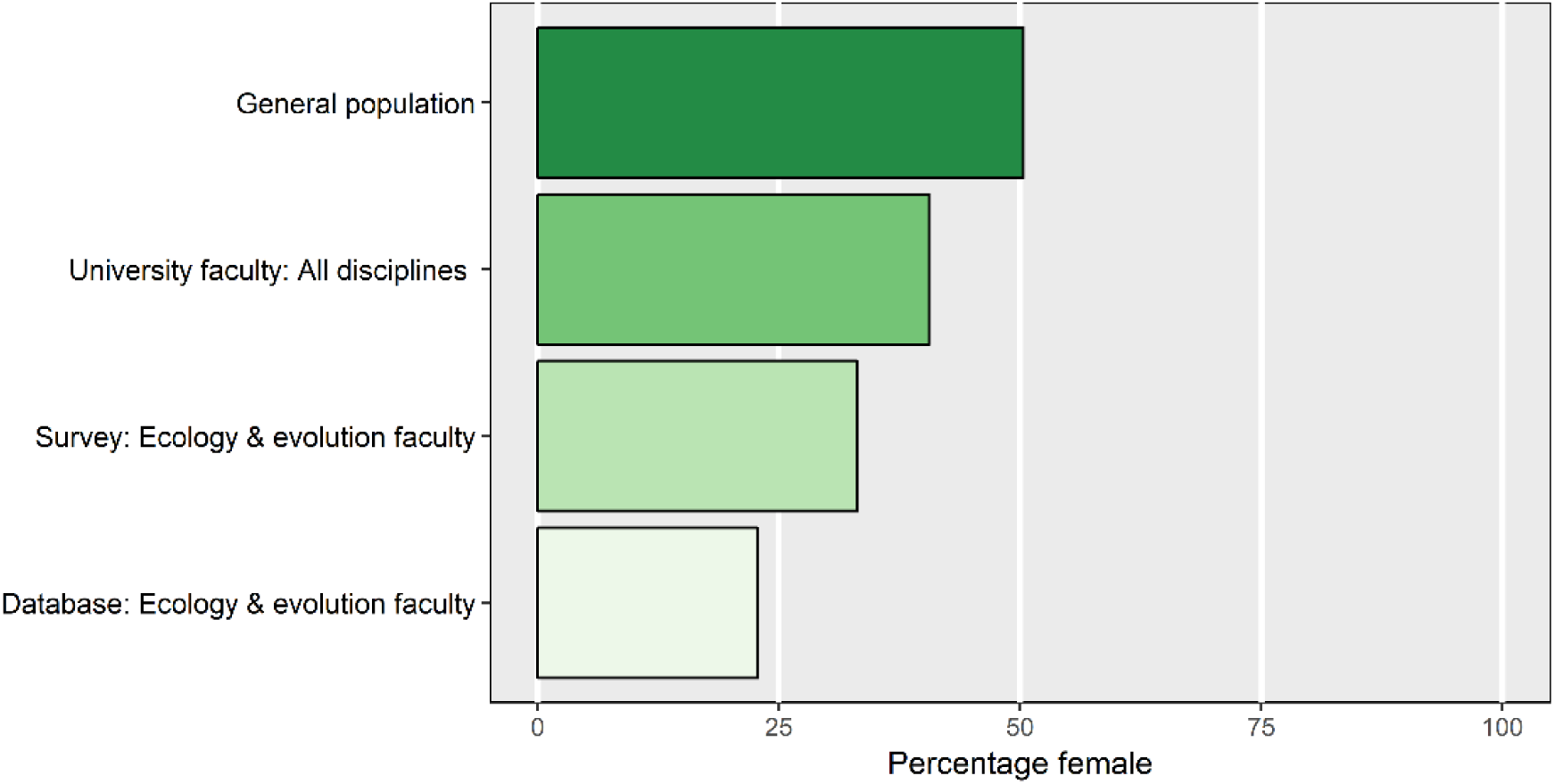
Female researchers are underrepresented, compared to the general population, among university faculty in general and ecology and evolution faculty. Source for general population and university faculty all disciplines: Statistics Canada. Source for ecology and evolution faculty: Researcher Survey and Researcher Database.

The percentage of female EE faculty varies between academic ranks, declining from Assistant Professor to Associate Professor to Full Professor. This pattern mirrors similar patterns in the general faculty population (Figure 3).

**Figure 3.**
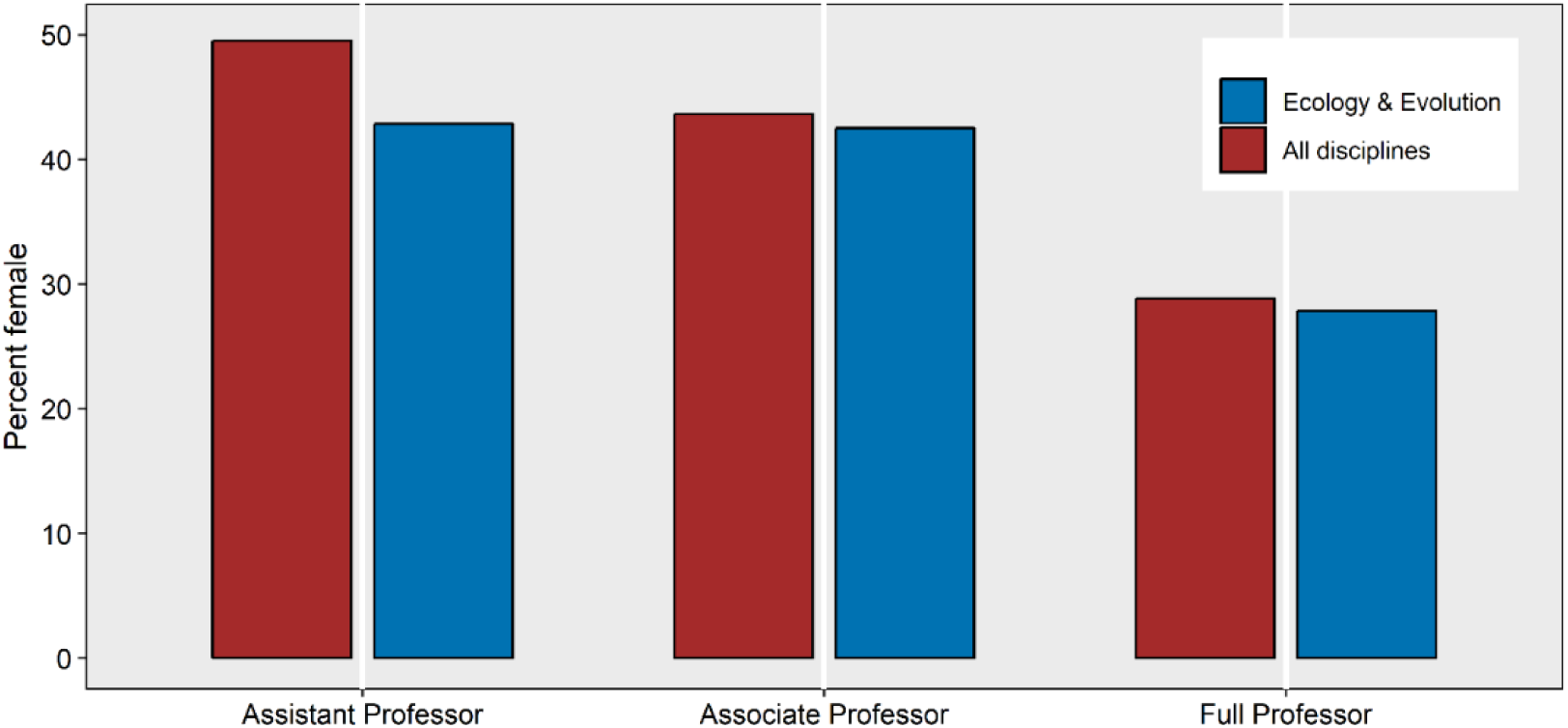
The percentage of faculty that are female declines from junior to senior academic ranks in both the EE and general faculty population. Source: Researcher database

Universities have increased the share of women among EE faculty hires. This can be visualized by examining cohorts of faculty who received their PhD in the same year: more recent cohorts are increasingly female (p < 10^−232^; Figure 4). For example, 28 researchers received their PhD degree in the year 1997, and 13 of them are female, so the percentage of female professors is 46%for the 1997 cohort.

**Figure 4.**
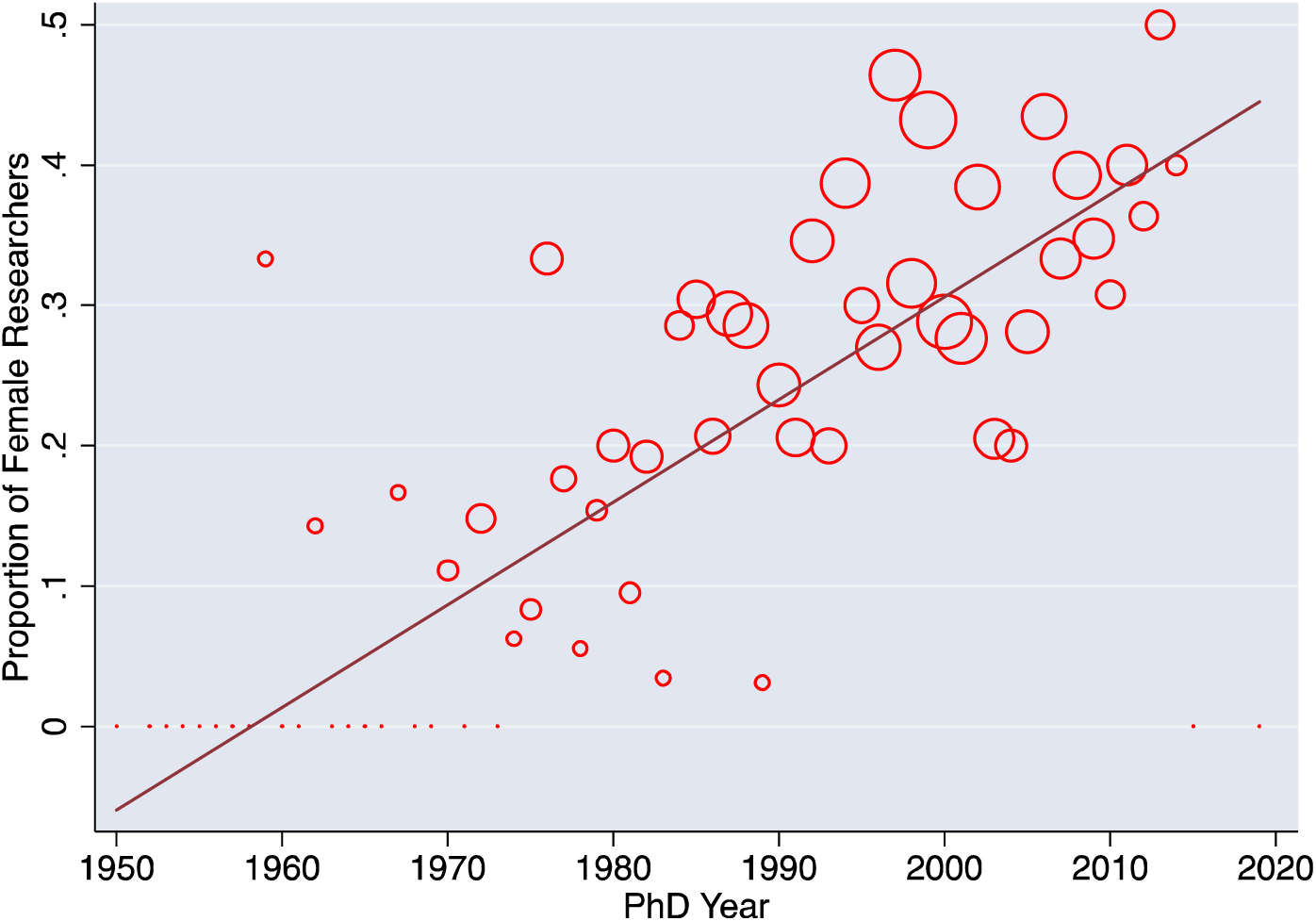
Female representation increases in cohorts of EE faculty (binned by PhD year) who received their PhD more recently. The size of the circle is the number of female faculty in the cohort. Source: Researcher database

In addition to gender, our survey also collected information about ethnicity of EE faculty (Figure 5). Indigenous researchers are dramatically underrepresented both in the overall faculty population and among EE faculty (1.2%). Visible minorities are also markedly underrepresented among ecology and evolution faculty (6.1%), compared not only to the general population but also to other disciplines in Canadian universities. The EE faculty population in Canada remains highly homogeneous in ethnicity.

**Figure 5.**
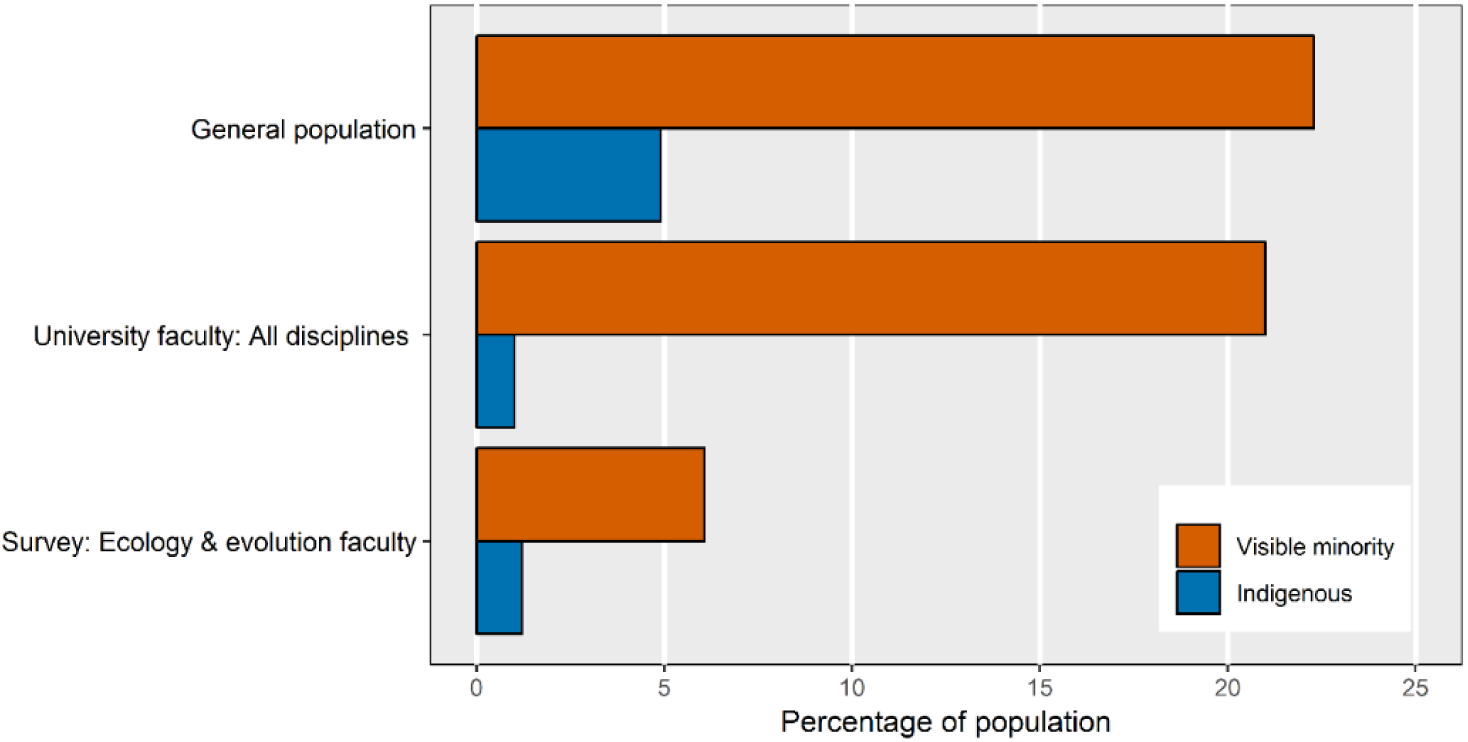
Visible minority and Indigenous researchers, compared to the general population, are underrepresented among EE faculty. Source: Researcher Survey (169 respondents).

### Participation in Working Groups, and Barriers to Inclusivity

The majority of the 169 faculty who took part in our survey had participated in at least one working group (Figure 6): 54%of female researchers, and 64%of male researchers (a non-significant difference: χ^2^ = 1.2, p =0.27).

**Figure 6.**
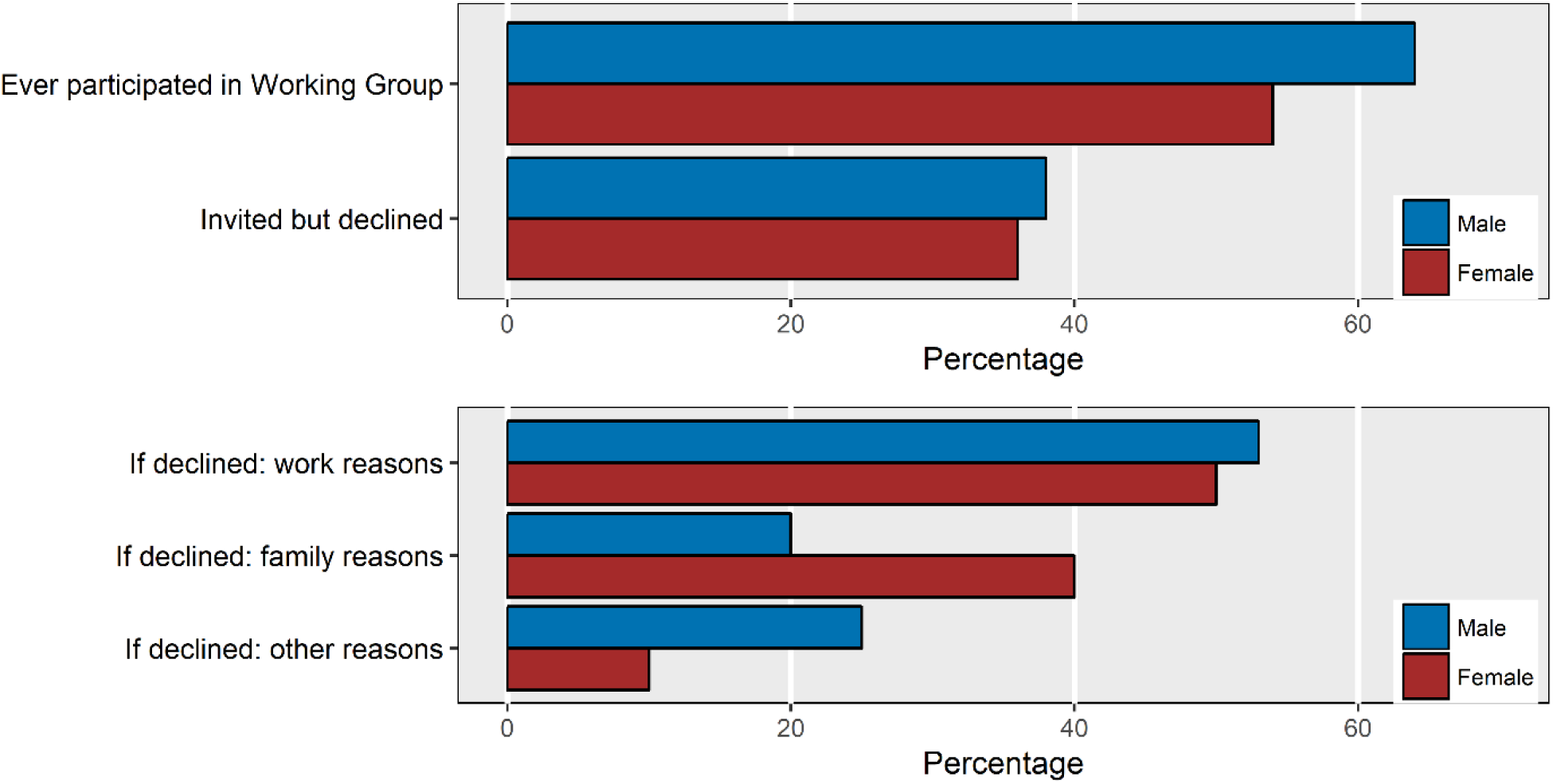
The percentage of faculty who have ever participated in a working group or have ever declined an invitation to participate are similar between male and female respondents (top panel, N = 169). The reasons for declining a working group invitation show more difference between male and female respondents (bottom panel, N = 63). Source: Researcher Survey

Our survey data provide more specific information regarding whether researchers have declined to participate in working groups and, if so, why (Figure 6). A similar proportion of female (36%) and male (38%) researchers declined at least one invitation to take part in a working group. However, the reasons for declining the invitation vary. Both genders list “work-related duties” as the main reason for declining an invitation, but the second-most common reason for female researchers is “family-related duties” whereas men are at least as likely to give other reasons, such as not being interested in the topic or not liking the working group method.

### Working Groups, Gender and Publication Impact

The first synthesis science centre, NCEAS (National Center for Ecological Analysis and Synthesis) was established in 1995 (Hampton and Parker, 2011). Since then, working groups have become an innovative and productive research approach used by more and more scientists. Canadian researchers have participated in working groups around the world, including those organised by the Canadian Institute of Ecology and Evolution (founded 2008). Among Canadian researchers, working group publications were evident soon after NCEAS was established in 1995, although for most researchers such working group publications account for a minority of publications each year (Figure 7). Using a fixed effects model, we find that there is no significant difference between male and female researchers in the proportion of their overall publications originating from working groups (interaction (gender * year): p = 0.47).This supports our earlier conclusion from the survey that male and female researchers have similar rates of participation in working groups.

**Figure 7.**
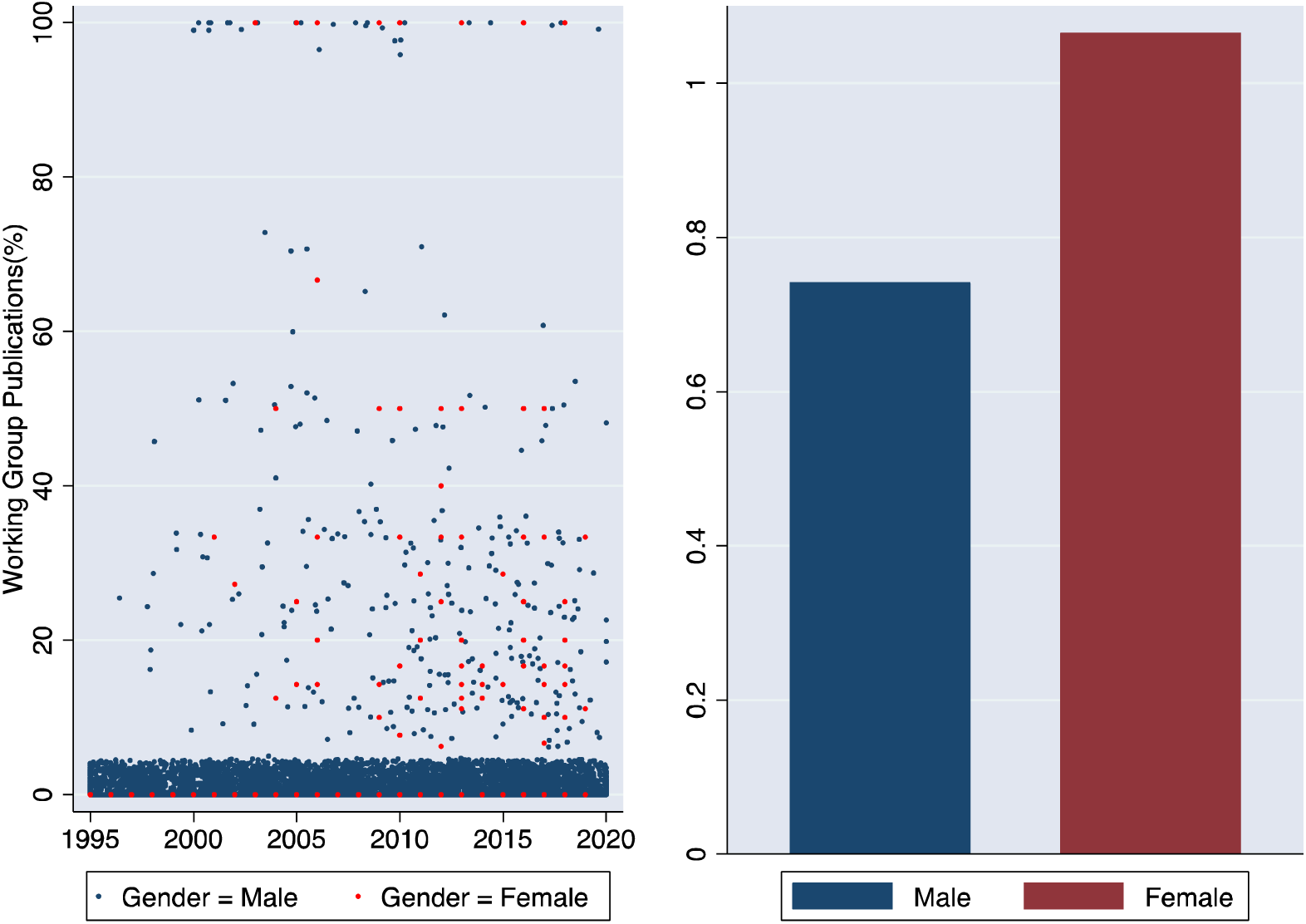
The percentage of working group publications of total publications in a particular year for both female and male researchers from 1995 to 2019, displayed either for all researchers (left panel) or as mean over both time and individuals within a gender (right panel). Note that the vertical scale on left and right panel differs, although the response variable is identical. Source: Researcher database

Our models explore the effects of working group participation and gender on the trajectory of H-indices over time. It should be noted that this is a longitudinal analysis, such that each researcher acts as their own control: the model compares the trajectory of their H-index before participating in a working group with that after the working group (although note that many researchers in our database never participated in a working group during our study period). Even so, we caution that it is difficult to unambiguously ascribe causality in observational data.

The first question we addressed was whether participating in working groups has a positive association with researchers’ H-indices. Based on our models, the answer is yes, but only later in the career (Figure 8; Model 2 in Table 1). For those who are fourteen or more years out from their PhD, a first working group experience is associated with a higher H-index. However, in the first 4-13 years post-PhD, H-indexes increase at a statistically indistinguishable rate before and after participation in a working group, and for researchers who obtained their PhDs within the last 4 years, H-indexes increase faster prior to working-group experience (Figure 8). The negative coefficient of main effect of working groups and the positive coefficient of interaction with time since PhD suggests a nuanced relationship: working group experience is predicted to have a negative effect on the H-indices of researchers at the early stage of their careers, but significantly improve senior researchers’ H-indices. Figure 9 illustrates this association of working groups and H-index.

**Table 1.** Two-way Fixed-Effects Regressions of H-index on Working Group Experience and Gender. Models 1-4 are based on the full researcher dataset, model 5 is restricted to researchers who received their PhD after 2000, and model 6 is restricted to researchers awarded a NSERC Discovery grant recently (2015 to 2019). Note: ^*^ p < 0.05, ^**^ p < 0.01, ^***^ p < 0.001

|  | (1) | (2) | (3) | (4) | (5) | (6) | (7) |
| --- | --- | --- | --- | --- | --- | --- | --- |
| Time since PhD | 1.80*** | 1.88*** | 2.01*** | 1.66*** | 2.04*** | 0.19 | 1.32*** |
| Working Group |  | -8.05*** |  | 8.39*** | -6.69*** | -2.06 | -6.08** |
| Working Group*Time since PhD |  | 2.40*** |  |  | 2.26*** | 1.12* | 2.05*** |
| Female*Time since PhD |  |  | -0.56*** |  | -0.44** | -0.54** | -0.61** |
| Working Group*Female |  |  |  | -6.54*** | -2.82* | -1.02 | -2.57 |
| Constant | 2.80** | 3.01*** | 3.95*** | 2.04* | 3.91*** | -1.23 | 0.74 |
| Individual FE | Yes | Yes | Yes | Yes | Yes | Yes | Yes |
| Year FE | Yes | Yes | Yes | Yes | Yes | Yes | Yes |
| N (Observations) | 35828 | 35828 | 35712 | 35712 | 35712 | 4551 | 19252 |
| N (Individuals) | 1247 | 1247 | 1244 | 1244 | 1244 | 324 | 772 |
| R<sup>2</sup> (within) | 0.79 | 0.81 | 0.79 | 0.80 | 0.81 | 0.85 | 0.83 |
| R<sup>2</sup> (between) | 0.081 | 0.0039 | 0.012 | 0.012 | 0.0021 | 0.015 | 0.060 |
| R<sup>2</sup> (overall) | 0.51 | 0.55 | 0.52 | 0.54 | 0.56 | 0.47 | 0.53 |

**Figure 8.**
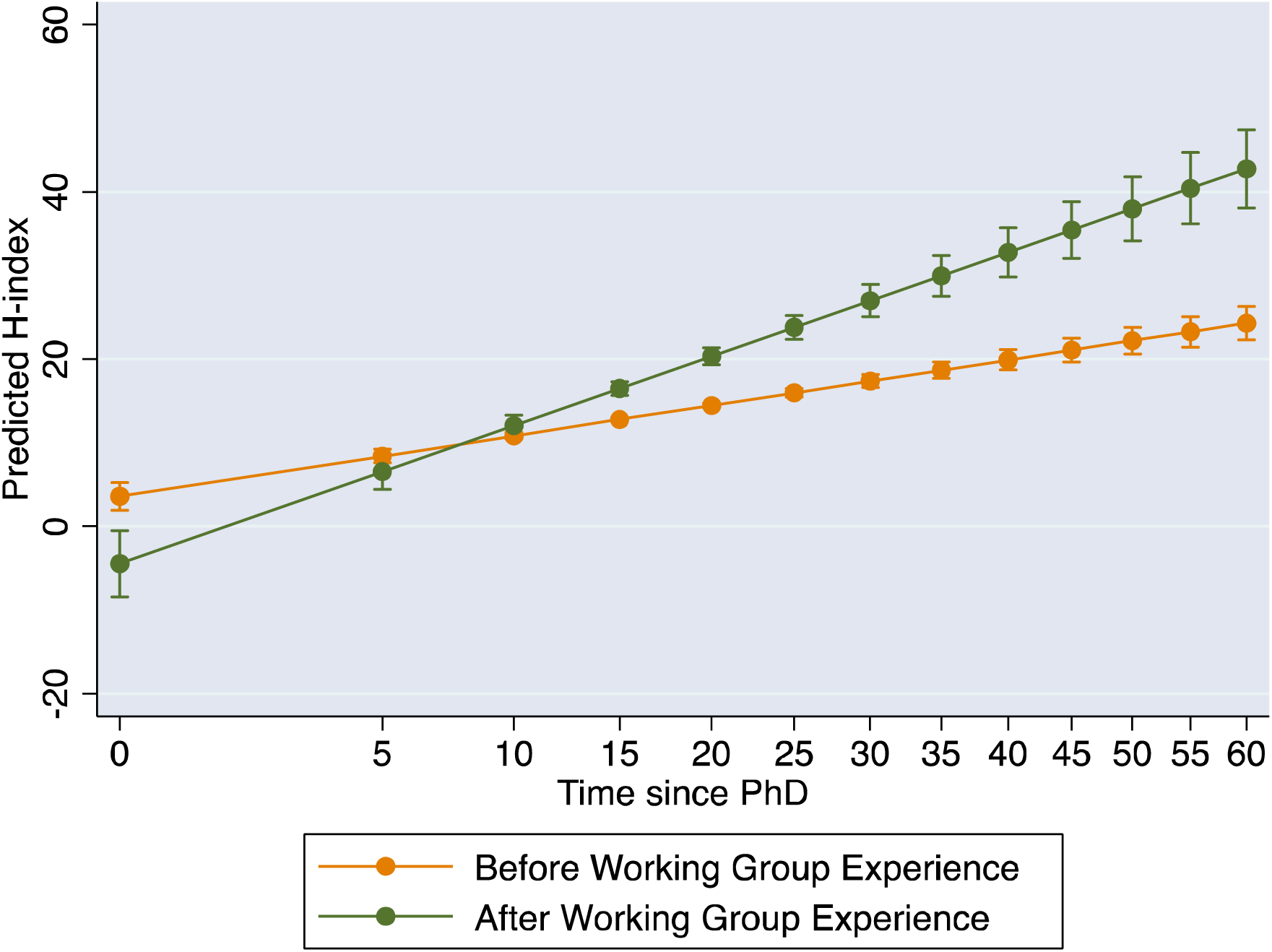
The predicted effects of working group experience on H-index are influenced by researcher gender and years since PhD. Error bars are 95% confidence intervals on predictions and the horizontal axis has a non-linear (year^0.6^) scale. Source: Researcher Database

**Figure 9.**
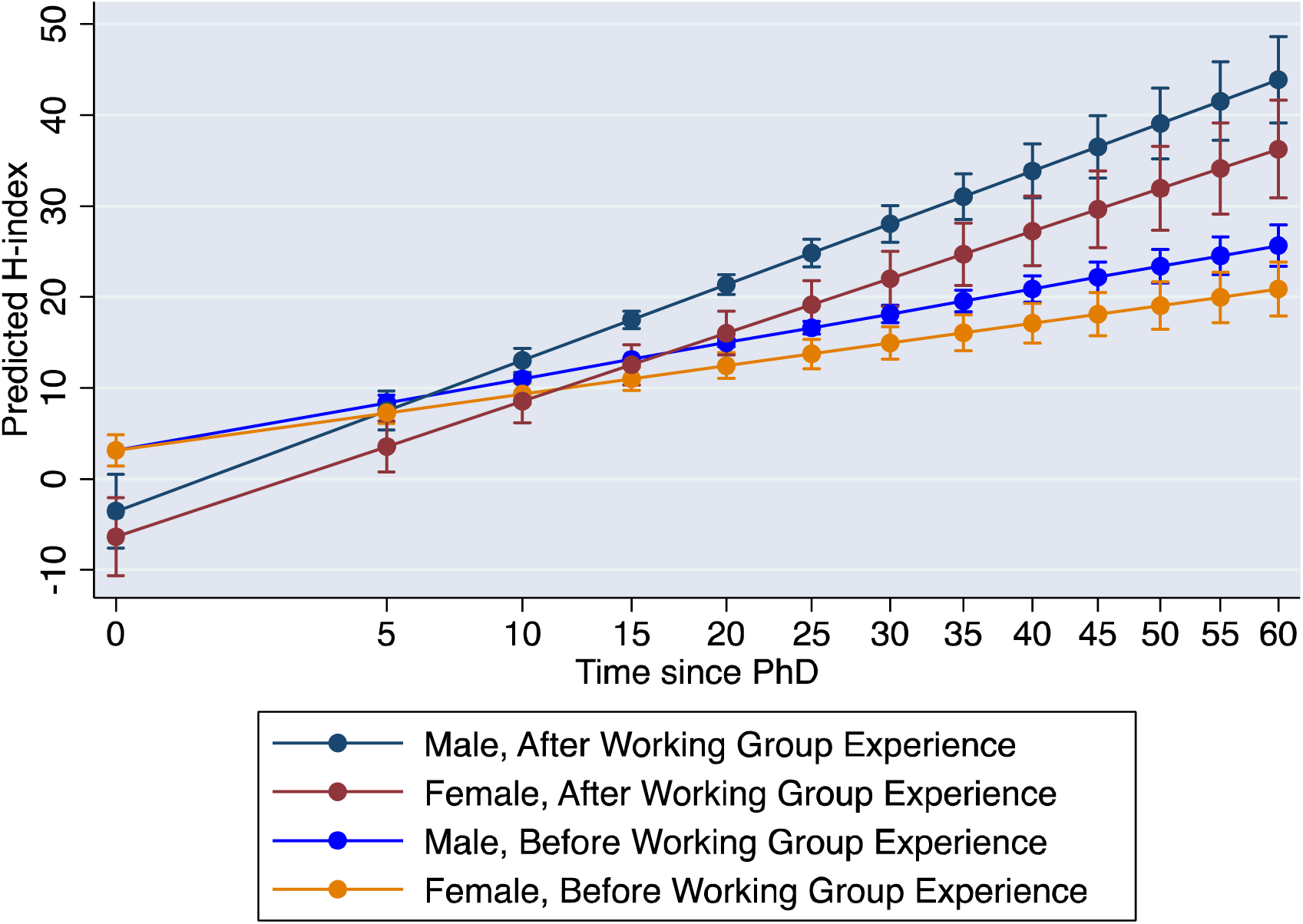
Predicted effects of working groups and gender on H-index. Error bars are 95% confidence intervals on predictions and the horizontal axis has a non-linear (year^0.6^) scale. Source: Researcher Database

Does this positive effect of working groups differ for male and female researchers? According to our findings, the answer is yes: in terms of advancing their H-index, male researchers benefit more from the experience of working groups than female researchers. Model 5 interacts gender with working group experience and also considers how the temporal change in H-indices is affected independently by gender and working group. Based on this model, we find male researchers benefit more from the experience of working groups than female researchers when it comes to advancing their H-index (Figure 9). We also considered a further three-way interaction with time, but the gender by working group effect does not significantly vary across career stages.

To facilitate visualizing the gender gap, we can examine the predictions separately for researchers with and without working group experience (right vs. left panels, Figure 10, based on Model 5). Regardless of working group experience, the gender gap - the difference between the H-index predictions for male and female researchers in each of these panels – is significant, but this gap widens with working group experience.

**Figure 10.**
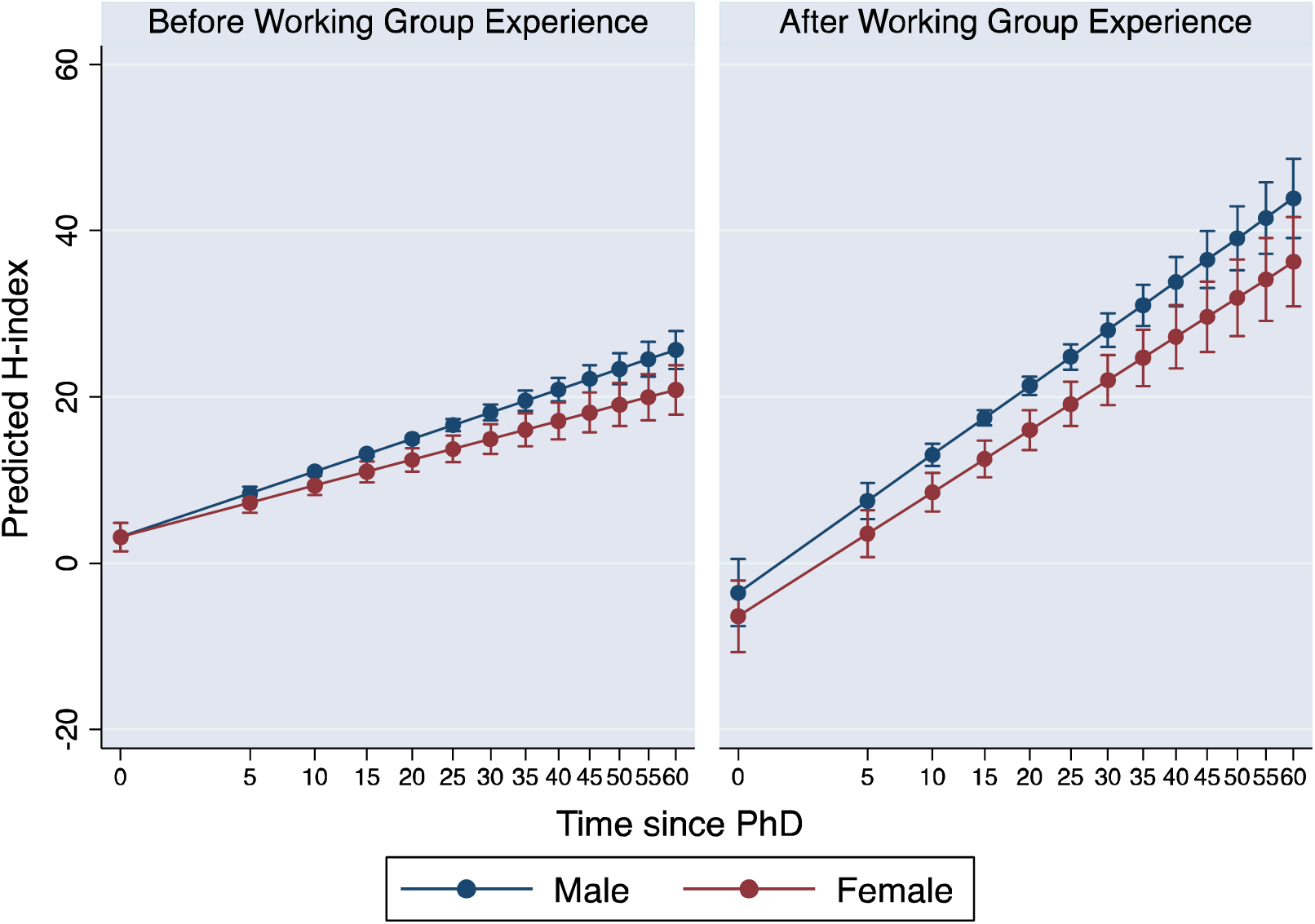
The gender gap in H-index increases with time regardless of working group experience, but is narrower for researchers without working group experience (left panel) than those without (right panel). Error bars are 95% confidence intervals on predictions and the horizontal axis has a non-linear (year^0.6^) scale. Source: Researcher Database

Finally, we ran two additional analyses focusing on different sub-groups. If we restrict our analysis to researchers who obtained their PhDs after 2000 (Model 6), the gender gap persists but is not exacerbated by working group participation (Figure 11). In other words, the gender gap in working group benefits is associated with cohorts of senior researchers. However, this younger cohort of researchers is still competing for grants against more senior cohorts of researchers. As mentioned above, our sample includes researchers who received their PhDs as early as the 1950s, who likely are no longer research-active or may even have died, so limiting our analysis to the subgroup of active faculty helps to establish a more accurate picture of the gender gap that today’s researchers face. If we consider currently research-active faculty, defined as all faculty who have won a NSERC Discovery grant during the past five years, from 2015 to 2019, the findings remain substantially the same as the full dataset: working group experience is associated with higher H-indices and this difference is gendered (Model 7). In practical terms, this means that the legacy of gender inequities from before the turn of the millennium may still be distorting the H-indices of today’s grant winners.

**Figure 11.**
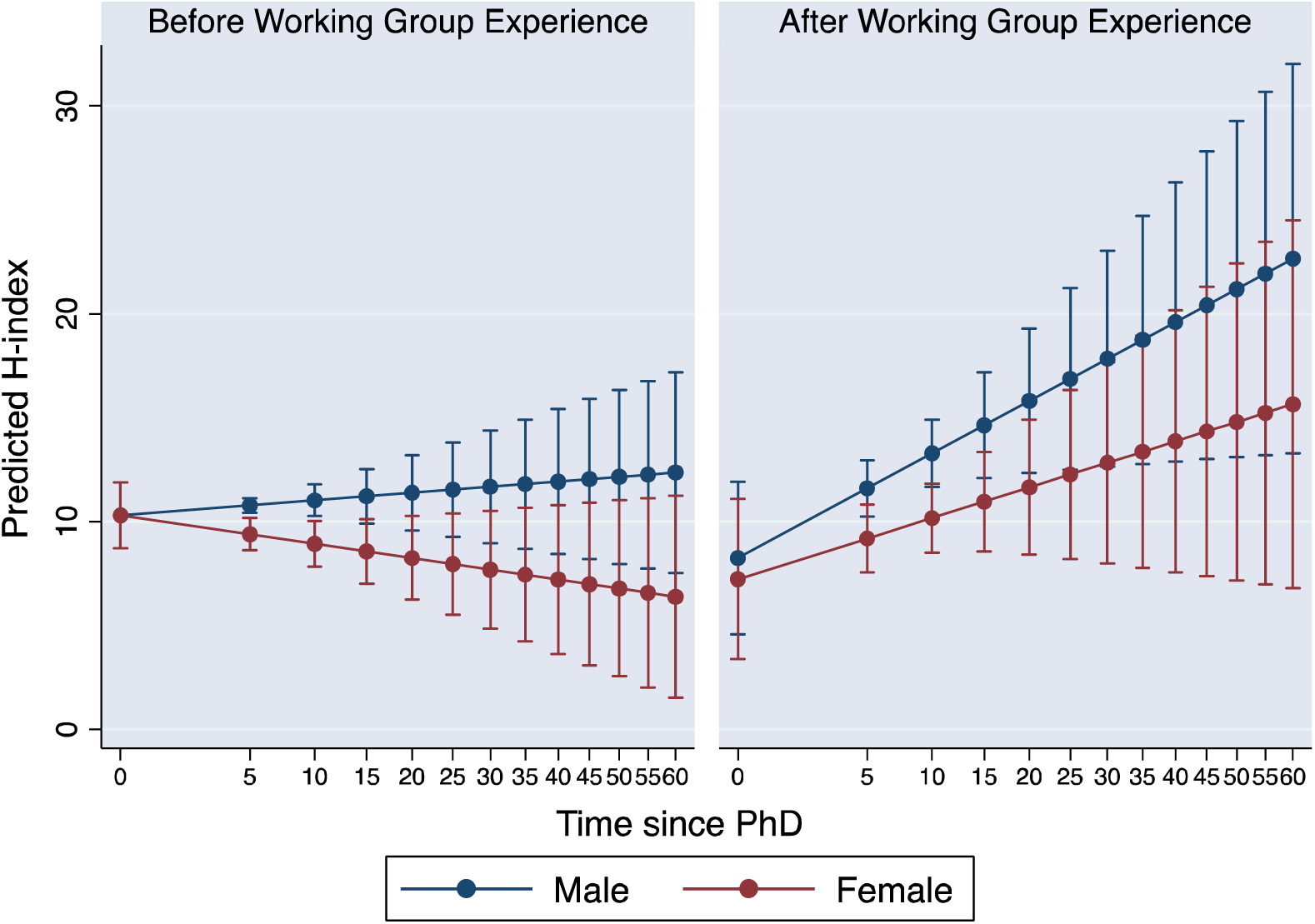
The gender gap in H-index for researchers who obtained their PhD less than 20 years ago (year 2000-2019). Error bars are 95% confidence intervals on predictions and the horizontal axis has a non-linear (year^0.6^) scale. Source: Researcher Database

### Potential mechanisms

Working groups may accelerate the H-indices of researchers in two different ways. First, if working group publications are more often cited than non-working group publications, participation in a working group could directly increase the number of highly-cited publications, the basis of the H-index. This could occur both because of the type of publication created by working groups - synthetic publications are cited more than primary research (Miranda and Garcia-Carpintero, 2018) - and because of the collaborative nature of working groups - publications from collaborations are cited more (Lariviere et al, 2015; Leimu and Koricheva, 2005). Second, working groups could provide future benefits to researchers, such as the development of collaborations after the working group, enhanced access to future funding opportunities, or re-use of databases developed by the working group in another project. For example, previous analyses of a U.S. synthesis centre found that its participants were more collaborative after participating in a working group (Hampton and Parker 2011). We found evidence for each of these effects in our study.

Working groups produce, overwhelmingly (>98%), synthesis science publications. Synthesis science publications, by themselves, are more cited than primary research publications (Figure 12; χ2 = 33.7, p = 10^−9^). Synthesis science publications can be divided into three types: mathematical (e.g. theoretical or simulation models), conceptual (e.g. literature reviews, framework papers) and statistical (e.g meta-analyses, species distribution models). All three types are cited more than primary research, but this is especially true of statistical syntheses (Figure 12; research type: χ^2^ = 177.5, p = 10^−16^). Independent of the type of synthesis research, publications from working groups are cited more than publications based on other, more traditional, methods (Figure 12; working groups: χ^2^ = 65.8, p = 10^−16^; working group x research type interaction, χ^2^ = 2.7, p = 0.44).

**Figure 12.**
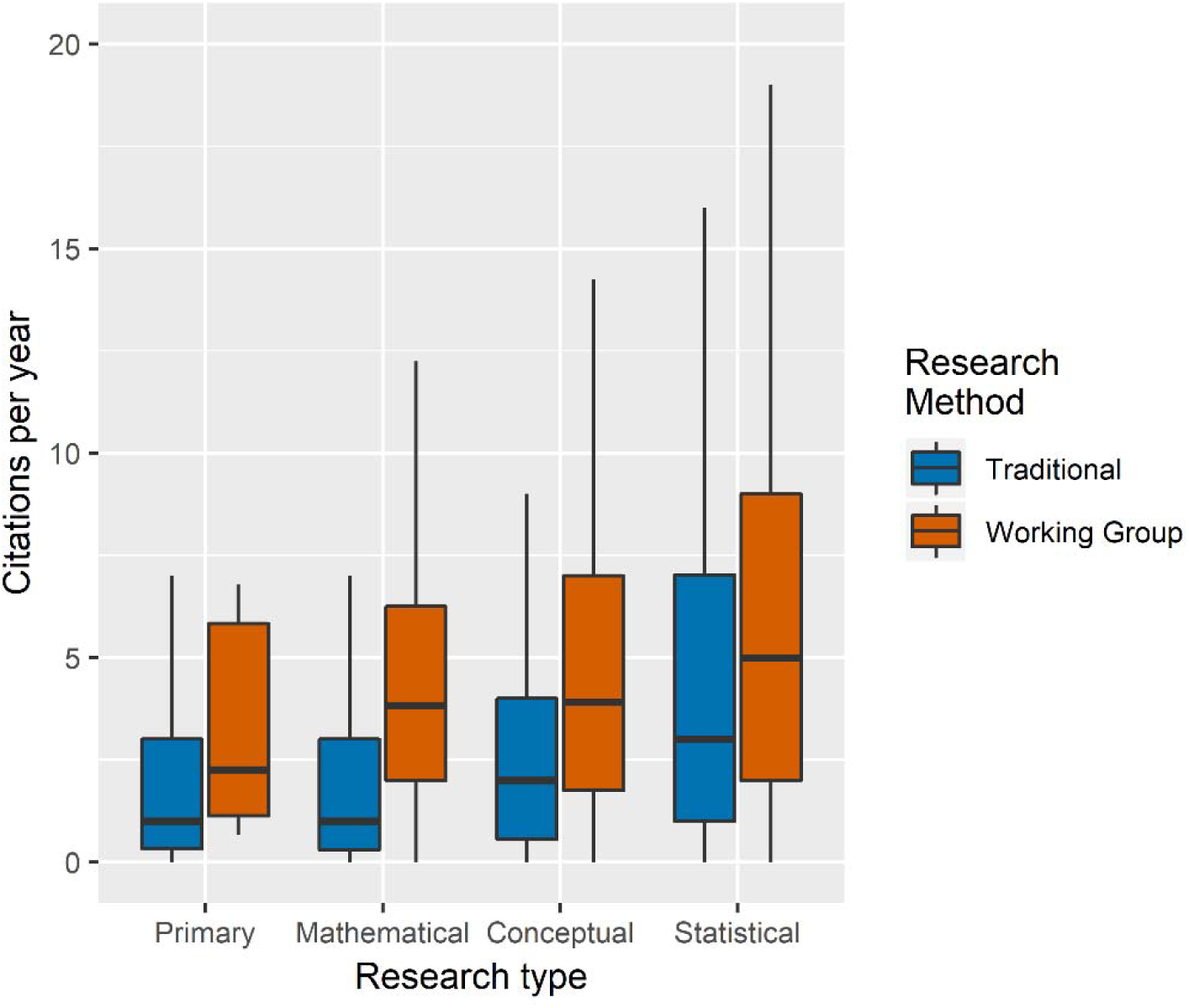
Publications vary in the average number of citations per year according to research type (primary research or synthesis research, with the latter comprising mathematical synthesis, conceptual synthesis and statistical synthesis) and research method (working group, or non-working group, here denoted “traditional”). Source: Publication database

Working groups also provide future benefits. In our survey, we asked the 102 researchers who had participated in a working group about indirect benefits of their most recent working group. The majority of respondents reported that they developed new collaborations in their working group which carried forward into new projects. Roughly a quarter of respondents reported that their participation in a working group led to new funding opportunities or the ability to reuse a database developed in a working group for a new purpose. Importantly, there was no gender difference in these proportions, suggesting that male and female researchers have similar future benefits, at least for their most recent working groups.

We plan further analyses of our dataset to partition the working group effect on H-indices into the direct effect of publications and the indirect future benefits, and to further establish the causal relationships.

**Figure 13.**
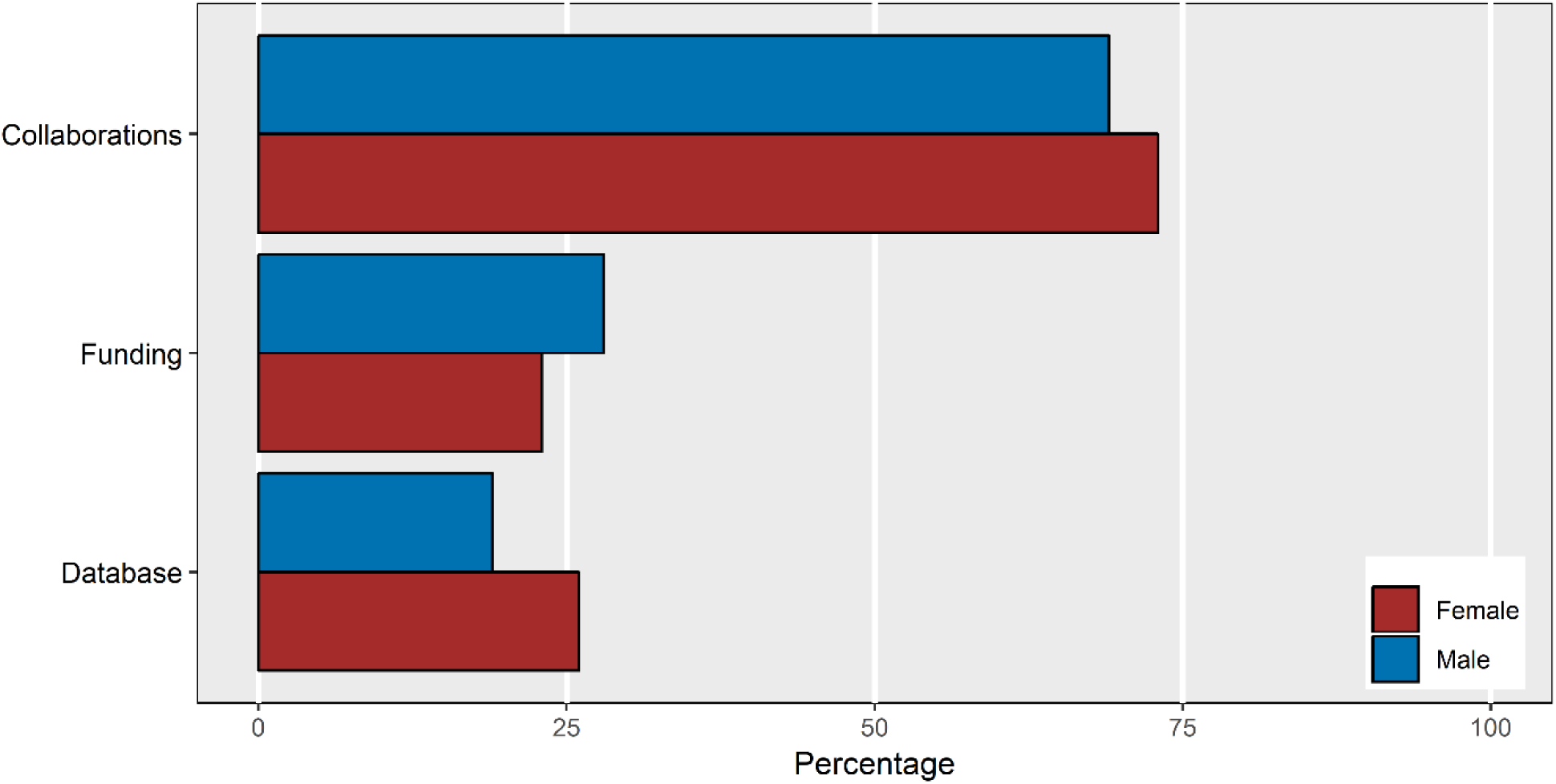
Percentage of researchers (N = 102) who received future benefits from their most recent working group in terms of subsequent collaboration with working group participants in other contexts (“collaborations”), funding opportunities (“funding”), and reuse of a database constructed within the working group for other projects (“database”). Source: Researcher database.

Our analysis also discovered that academic age and gender moderated the effects of working group experience on H-indices. There are several potential mechanisms here.

First, this may represent real differences between working group participants in the costs born and the benefits reaped. The publications produced by working groups are the result of many different activities, such as: collating of large amounts of published data into databases, qualitative summaries of the literature, advanced statistical analyses or mathematical modelling, creation of figures, and writing and editing of manuscripts. These activities may not be shared equitably. When graduate students and postdoctoral fellows are relied upon to do the more time-consuming and routine tasks (often collating data and surveying literature), they may have less time to invest in primary research. Previous studies suggest that scientific labour in research collaborations is gendered, with women more likely to collect data and men more likely to make conceptual, material, statistical and editorial contributions (Macaluso et al. 2016). The benefits may not also be shared equally. Because publications tend to increase the reputation of first authors or last authors more than other co-authors, if such authorship positions are disproportionately claimed by older and/or male researchers (West et al. 2013) then less benefit from publications will accrue to other researchers.

Second, there is a historical dimension to patterns in researchers’ H-indices. The H-index is a cumulative measure of publication impact, and so it preserves - over the entire career of a researcher - any historical inequities in the distribution of publications or citations in the researcher community. Since the citation rate of papers increases with time since publication, the H-index may in fact magnify such historic patterns. Further analyses of our dataset will enable us to more fully characterize how the impact of academic age, gender and working group participation may have changed through time.

Many synthesis centers have enacted policies to ensure that the participant composition is gender-balanced and represents a range of academic career stages and backgrounds. Such policies probably explain why male and female researchers in our survey reported similar rates of participation in working groups. The challenge is to now go beyond these current policies to ensure that the career benefits of participating in working groups are equitably shared amongst participants.

## Acknowledgements

This project was funded by a NSERC “Studies in NSE Research in Canada” grant to DSS, SF, and C C-B. The survey was conducted under Human Ethics certificate H19-00891 approved by the University of British Columbia’s Behavioural Research Ethics Board. We thank the undergraduate assistants who helped build the databases - Mirkka Puente, Rodrigo Vallejo, Liam DelGesso – and Kelly Haller for assistance with administration, survey logistics and reporting.

## Appendix 1. Additional information included in the Researcher Database

### Academic career stages

We recorded the start and end of postdoctoral fellowships, appointment as an assistant professor, and - as relevant - year of tenure and promotion to associate professor, year of promotion to full professor, and year of appointment as emeritus professor. This information was largely obtained from online CVs and researcher-maintained websites. Where necessary, we made some assumptions about the career progression of researchers. We assumed that the tenure track period began with appointment as an Assistant Professor, or its french equivalent *professeur adjoint*, and that tenure was granted at the time of promotion to Associate Professor, or its french equivalent *professeur agrégé*. Although tenure and promotion are decided separately, they usually coincide in Canadian universities. If only the year of tenure and the year of promotion to Full Professor were known, we inferred that the researcher was an Associate Professor in the intervening period. We did not assume that all retiring professors became emeritus, instead we searched web resources for the title of Emeritus Professor.

### Administrative appointments and years

We also collected information on administrative appointments, including the year started, year ended and position (head, dean, vice president, assistant dean and director). This information was relevant to the study as faculty may be less research-active during administrative appointments.

### Adjunct professors

Adjunct Professors often have a primary employer outside of the university, such as a governmental agency or private company. The career paths of Adjunct Professors are varied, and some differ substantially from tenure-track university appointments. We identified all researchers that were Adjunct Professors by searching for this title or its French equivalent, *professeur associé*.

## Appendix 1 WOS Data Cleaning & Record Linkage

Although there were important variations in the Python record linkage script we used, in the description below, we explain the umbrella strategy that we mobilized to generate a clean publication record.

### Filter I - Publication Names

The general goal of record linkage in our case is to remove publications that do not match a set of unique identifiers associated or attached to our researchers. For the first filter, we used the name(s) used by the focal scholar in their publications. Therefore, for each case, the first step is to filter out the WOS publications for which the author(s) fields do not contain the “known publication names” of our scholar. For scholars with GSC-level data, the Python program automatically generates a list of all the spelling/naming variations/conventions of the names as they appear in the bibliometric data-points. Most researchers have one consistent name, but enough have a second variant that the mean name variants overall is 1.85. For all the other cases, our team of research assistants manually generated the “publication name(s)” list. We implemented this filtering step as a Boolean test; either the known name was in the author column or not. On average, this first filtering step removed 14%of publications contained in the raw csv file.

### Filter II - Institutional Affiliations

Working with the remaining publications, the second filtering step involved matching known and validated institutional affiliations (graduate schools, early career & current positions) that we collected for each scholar both programmatically and manually from a variety of sources (CVs, NSERC api, departmental, personal, and/or lab websites) with the list of authors’ institutional affiliations in the WOS meta-data. On average, we collected three different institutional affiliations. In this filtering test, for each publication remaining from Filter I, if the Python program successfully matched any of those school names to the list of institutional affiliations for the author with the same name, then the publication was kept in the corpus. Here, after some string cleaning, as with the first filtering step, the Python Program implements a Boolean test. In other words, the result of the matching test between every known school name and WOS institutional affiliation was a binary output: match or no match. On average, this second filtering step is by far the most efficient as it shaves off 41%of all WOS publications remaining after the first filter (name) was applied.

### Filter III - Publication Title Matching with Fuzzy Logic

Although filter I and II might seem robust enough, they do not constitute record linkage properly speaking. However, linking publication titles from CV and/or GSC profile with WOS record is and it constitutes a robust validity test. Therefore, for cases with CV and/or GCS-level data, the Python program uses fuzzy matching on the corpus of remaining WOS publications to compare article titles. Fuzzy logic is a form of multi-valued logic that deals with a reasoning that is approximate rather than fixed and exact. Fuzzy logic values can range between 0 and 1. In contrast, Boolean logic –as used for Filter I and II–Is a two-valued logic; True or False. When comparing two strings using fuzzy logic, we are trying to answer the question “how similar are string A and string B?”. Boolean logic, on the other hand, asks “are string A and string B the same?”. When using fuzzy logic for string matching, we normally talk of fuzzy string matching. It is a process of finding strings that approximately match a pattern. The process has various applications such as spell-checking, DNA analysis, spam email detection, and plagiarism detection (Zadeh, 1988). In our Python program we use the package “Fuzzywuzzy” which relies on Levenshtein Distance to calculate the difference between given sequences and patterns.

In our Python code, each WOS title remaining after the second filtering step is compared to all the publication titles from the CV/GSC corpus. All matches above 0.75 are kept in an array and at the end of the loop the WOS title with the highest fuzzy score is flagged as a match. As with other rules-based computational approaches, here the cut-off point of 0.75 was generated manually after carrying out thousands of benchmarking tests and manually analyzing the outputs with descriptive statistics. If no title’s fuzzy score reached the cut-off of 0.75, then the WOS title is considered a non-match. On average, this third filtering step removed 26%of the publications remaining in the corpus of articles potentially constituting the publication record of our scholar.

The corpus of WOS publications remaining after this triple filtering process constitutes the base of each scholar’s publication record. On average, 67%of all publications in the final csv of our sampled population originated from this primary matching/record linkage algorithm (primary corpus). One-third comes from the secondary cleaning program which we explain next.

### Filter IV - Secondary Cleaning with Additional Institutional Affiliations

A rule-based computer program like the one we develop is not necessarily static as it can also be generative of new information that can in turn be used recursively to match additional publications previously “missed” or filtered out. That is, information not known at run-time. The Python program does that in two ways.

First, we take advantage of the specific way that WOS curated the authors’ institutional affiliations meta-data. Since this column can contain more than one institutional affiliation per author (i.e., [Adam Zhang, Univ Toronto], [Adam Zhang, Univ Alberta]), the Python program is able to identify new institutional affiliations previously unknown to us. We only ran this code on the primary corpus, namely articles that successfully passed the first three filtering rounds. To generate “new schools”, the Python program iterates through every publication in the primary corpus and flags any institutional affiliation(s) attached to our scholars that are not in our list of known institutions. This is an extremely reliable way of generating new information since we are only running this code on publications with matched name, institution, and title. Manual inspection of cases uncovered with “new schools” reveals that those institutional affiliations tend to be short-term: post-doctoral fellowship, visiting position, government-run organizations, etc. On average this method yields 0.87 new institutional affiliations per researcher.

In terms of programmatic steps, if at least one “new school” is discovered with this protocol, the Python program will (1) review all the cases that were dropped between filter I and filter II, (2) identify those containing any “new school(s)”, (3) recursively apply filter III to these cases, and (4) finally update the primary corpus if need be. That said, those familiar with the concept of recursive function in computer science (Anderson, Pirolli, and Farrell, 1996) will notice a hidden fifth step here. Namely, if at least one new publication is found, the Python program will re-do steps (1) to (4) until no new case is found.

The second source of additional institutional affiliation(s) comes indirectly from GSC. The Python program first generated a secondary corpus composed of all the WOS publications that passed filter I and III (name and GSC titles), but failed filter II (institutional affiliation). Then the algorithm yields, as explained above, a list of previously unknown schools. Finally, the Python program is recursively re-run with that new information and the new matched publications are vetted before being included in the final corpus. On average this method yields 1.70 new institutional affiliations per researcher. Filter IV accounts for one-third of all publications in the final corpus.

## Appendix 2 Identification of Working Group Publications

### Method 1- Fuzzy matching of publication titles from Synthesis Centre records vs. WOS publication record

As the first method to identify synthesis research in our curated WOS publication corpus, we use the same fuzzy logic algorithm developed for the WOS filtering process (Filter III) to match synthesis centres’ publication titles with titles from our sampled populations’ WOS publication record. When the publications listing from synthesis canters included not only titles but the full references, we used a Ruby programming language Gem called “AnyStyle” to parse the title from the bibliometric data. After manually controlling for false positives, we matched 512 WOS titles from 165 different researchers’ publication record with the corpus of synthesis centres’ publication titles (11%).

### Method 2 - WOS keywords matching synthesis centres names and working groups descriptors

As a second method to identify synthesis research titles in the cleaned WOS corpus of publications from our sampled population of ecology and evolution researchers, our team generated a list of 39 keywords (see Table 2) covering both known synthesis centres and 6 keywords that represented common descriptors of working group (“working group”, “synthesis group”, “synthesis working group”, “synthesis committee”, “synthesis workshop”,”catalysis group”). We ran a Python Program that generated an output containing all the WOS publications for which at least one of the keywords was found in one or several of the six metadata columns from the WOS csv (title, abstract, abstract keywords, funding, acknowledgments, WOS keywords). This keyword matching effort yielded 368 new WOS titles.

**Table 2:**
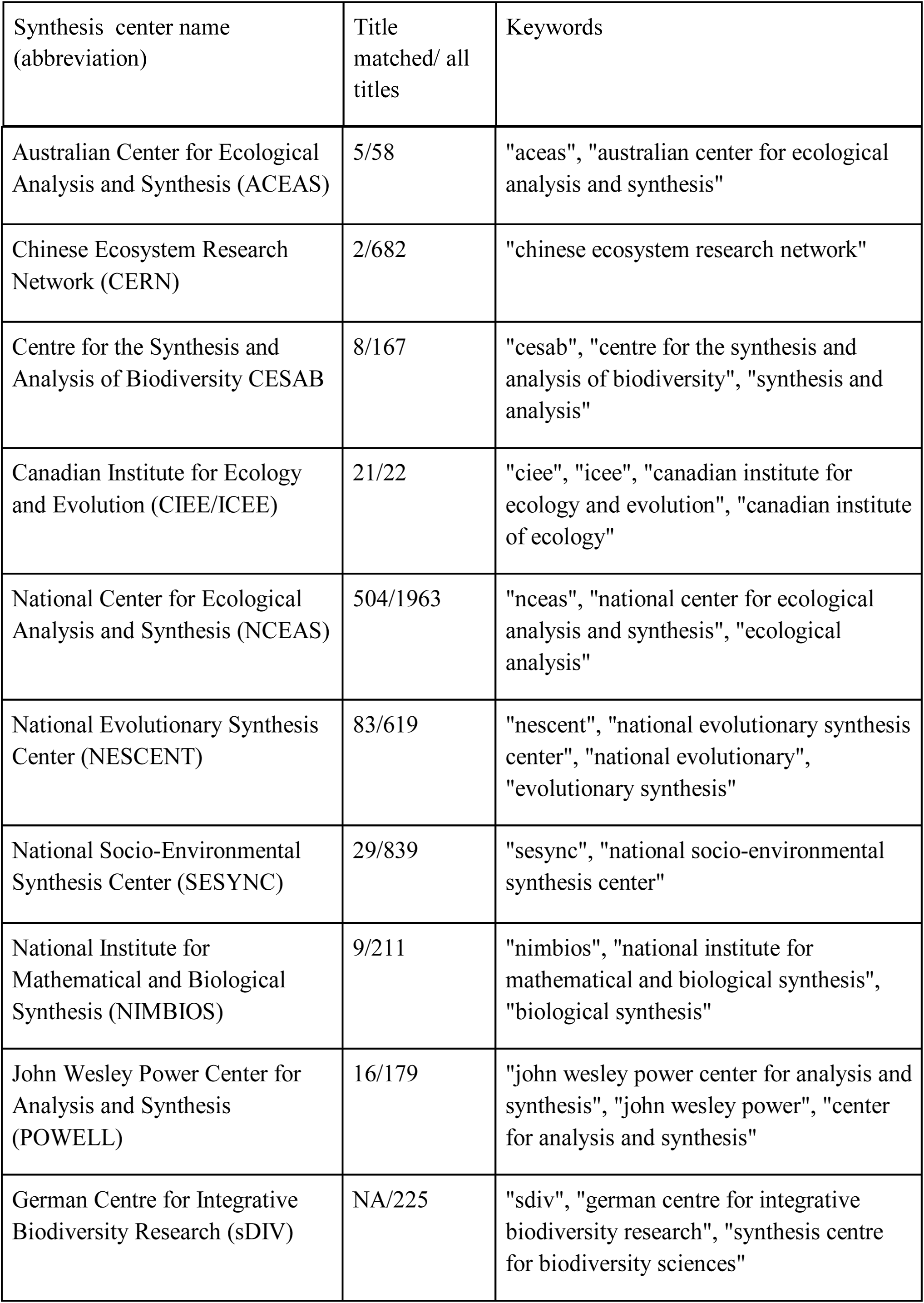
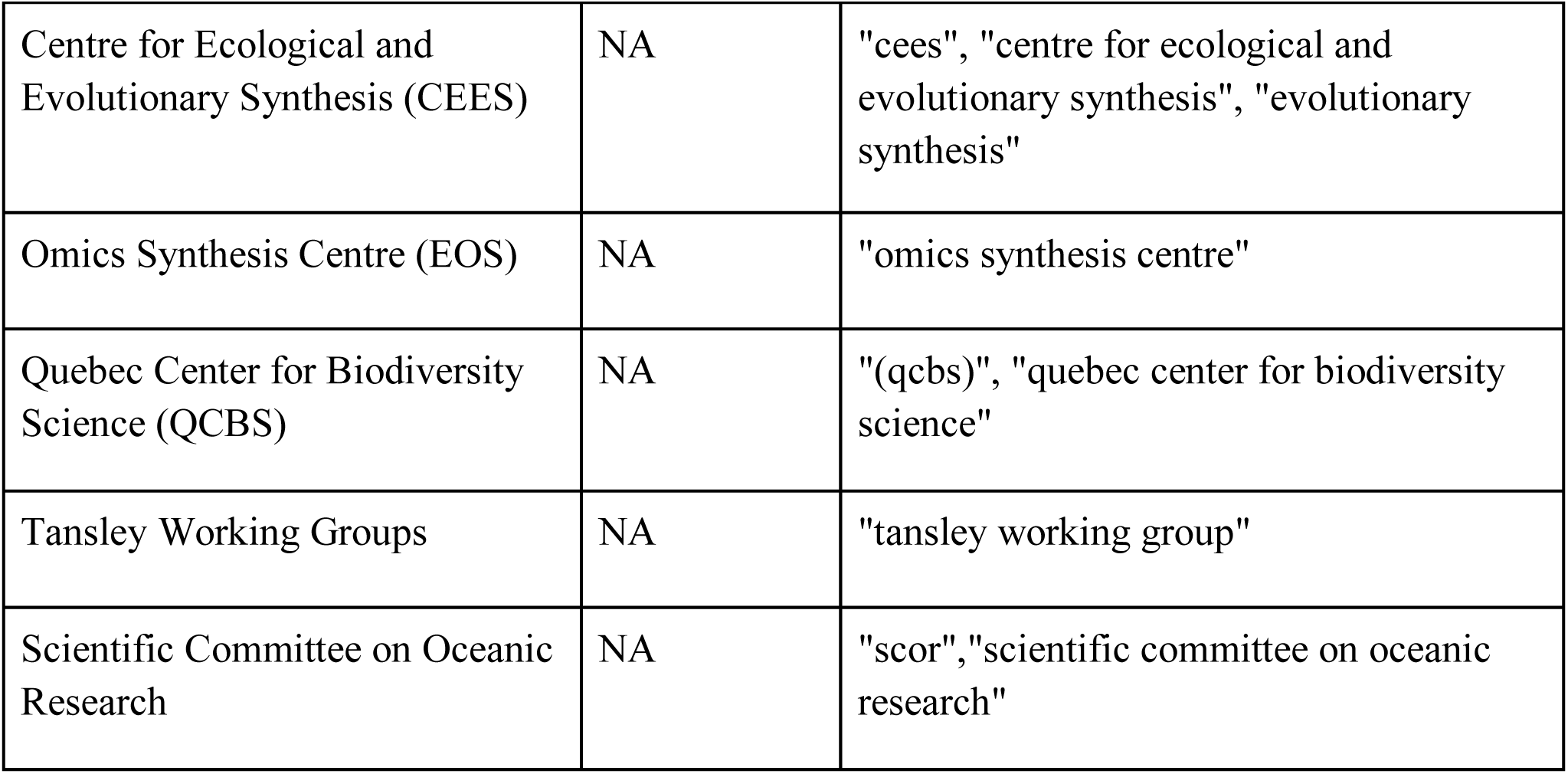
Synthesis centres used to search for working group publications in our database, either by comparison of WOS publication titles with the list of publication titles available from each centre (NA = Method 1 not used for that particular centre) or by searching for keywords that represented the centre.

| Synthesis center name<br>(abbreviation) | Title<br>matched/ all<br>titles | Keywords |
| --- | --- | --- |
| Australian Center for Ecological Analysis and Synthesis (ACEAS) | 5/58 | "aceas", "australian center for ecological analysis and synthesis" |
| Chinese Ecosystem Research Network (CERN) | 2/682 | "chinese ecosystem research network" |
| Centre for the Synthesis and Analysis of Biodiversity CESAB | 8/167 | "cesab", "centre for the synthesis and analysis of biodiversity", "synthesis and analysis" |
| Canadian Institute for Ecology and Evolution (CIEE/ICEE) | 21/22 | "ciee", "icee", "canadian institute for ecology and evolution", "canadian institute of ecology" |
| National Center for Ecological Analysis and Synthesis (NCEAS) | 504/1963 | "nceas", "national center for ecological analysis and synthesis", "ecological analysis" |
| National Evolutionary Synthesis Center (NESCENT) | 83/619 | "nescent", "national evolutionary synthesis center", "national evolutionary", "evolutionary synthesis" |
| National Socio-Environmental Synthesis Center (SESYNC) | 29/839 | "sesync", "national socio-environmental synthesis center" |
| National Institute for Mathematical and Biological Synthesis (NIMBIOS) | 9/211 | "nimbios", "national institute for mathematical and biological synthesis", "biological synthesis" |
| John Wesley Power Center for Analysis and Synthesis (POWELL) | 16/179 | "john wesley power center for analysis and synthesis", "john wesley power", "center for analysis and synthesis" |
| German Centre for Integrative Biodiversity Research (sDIV) | NA/225 | "sdiv", "german centre for integrative biodiversity research", "synthesis centre for biodiversity sciences" |
| Centre for Ecological and Evolutionary Synthesis (CEES) | NA | "cees", "centre for ecological and evolutionary synthesis", "evolutionary synthesis" |
| Omics Synthesis Centre (EOS) | NA | "omics synthesis centre" |
| Quebec Center for Biodiversity Science (QCBS) | NA | "(qcbs)", "quebec center for biodiversity science" |
| Tansley Working Groups | NA | "tansley working group" |
| Scientific Committee on Oceanic Research | NA | "scor", "scientific committee on oceanic research" |

## Appendix 3

**Table 3:** Three Types of Synthesis Research. List of keywords used to identify additional synthesis research publications in the WOS corpus (Method 3).

| **Meta-Analysis**<br>**Keywords** | **Systematic Review**<br>**Keywords** | **Mathematical**<br>**Modelling** |
| --- | --- | --- |
| "meta-analyses" | "conceptual framework" | "individual-based model" |
| "meta-analytic" | "conceptual synthesis" | "mathematical model" |
| "meta-analytical" | "literature review" | "simulation model" |
| "metaanalysis" | "literature synthesis" | "stochastic model" |
| "meta-study" | "qualitative synthesis" | "theoretical framework" |
| "meta-studies" | "systematic review" | "theoretical model" |
| "meta-summary" |  |  |
| "metasynthesis" |  |  |
| "meta-synthesis" |  |  |
| "quantitative synthesis" |  |  |
| "vote-counting" |  |  |

## Appendix 4. Online survey

The following is the text of the survey distributed to faculty members as part of this study:

Thank you for agreeing to participate in this survey. We are interested in understanding the role of synthesis science – especially that conducted in “working groups” – in the ecology and evolution research community. We anticipate that this survey will take 5-10 minutes.

### PART 1: Background

*First, we would like to ask a few background questions.*

1. What is your name (format: first name last name)
2. What is your current academic status: Graduate student; postdoc; assistant professor, associate professor; professor; retired professor; non-academic researcher; other] [if **professor, then go to the following academic progression module]
  - Year completed PhD:___
  - Year first appointed to a tenure track position as Assistant Professor: ___
  - Year first achieved tenure: ___
  - Year first promoted from Associate to Full: ___
  - Year retired: ___
  - Please enter senior admin positions (e.g. head, director, assistant dean, deans, VP) and years
  - Head, from ___ (Year) to ___ (Year)
  - Director, from ___ (Year) to ___ (Year)
  - Assistant Dean, from ___ (Year) to ___ (Year)
  - Dean, from ___ (Year) to ___ (Year)
  - VP, from ___ (Year) to ___ (Year)
  - Provost, from ___ (Year) to ___ (Year)
  - Other, please specify ___, from ___ (Year) to ___ (Year)
  *Family demands can impact researchers’ trajectories, so we would like to start by asking you a bit about your family circumstances*
3. Do you have children?
  □ Yes
  □ No
  - *[If no, route to question 4]*
  - If yes, how many children do you have*? [set up next questions based on number of children]*
  □ 1
  □ 2
  □ 3
  □ 4
  □ 5
  □ 6 or more
  - In what year was your first child born?
  - In what year was your second child born?
  - In what year was your third child born?
  - ….
  - Have you taken parental leave?
  - *[If no, route to question 4]*
  - If yes, approximately how many months of parental leave have you taken, in total

> Child 1 From ___/___ (Month/ Year) to ___/___ (Month/Year)
>
> Child 2 From ___/___ (Month/ Year) to ___/___ (Month/Year)
>
> Child 3 From ___/___ (Month/ Year) to ___/___ (Month/Year)
>
> Child 4 From ___/___ (Month/ Year) to ___/___ (Month/Year)
>
> Child 5 From ___/___ (Month/ Year) to ___/___ (Month/Year)
>
> Child 6 From ___/___ (Month/ Year) to ___/___ (Month/Year)
>
> Add here if you have 7 or more children
  *We are interested in the degree to which synthesis centres promote equal opportunity for everyone, so we would like to ask you about your visible minority status*
4. In terms of ethnicity, do you identify as (Please check all that apply):

Canadian Aboriginal (First Nations, Inuk, Metis)

White

South Asian (e.g. East Indian, Pakistani, Sri Lankan, etc.)

Chinese

Black

Filipino

Latin American

Arab

Southeast Asian (e.g. Vietnamese, Cambodian, Laotian, Thai, etc.)

West Asian (e.g. Iranian, Afghan, etc.)

Korean

Japanese

Other - specify

### PART II: Synthesis Science in Working Groups

We would like to ask you some questions about your participation doing **Synthesis Science in Working Groups**. A working group is defined as a group of people that meets together for a period of intense collaboration, normally 3 to 5 days, to either develop new theory or to collate and analyse previously collected data. Working groups are often hosted and funded by synthesis centres, but may also be funded by other institutions, networks or granting agencies.

1. Are you currently or have you ever been part of a Working Group?
  - *[If no, route to last question]*
  - If yes, how many different working groups have you participated in? Many working groups have more than one meeting, please enter the number of distinct groups irrespective of the number of meetings *[set up next questions based on number of workgroups participated in]*
    □ 1
    □ 2
    □ 3
    □ 4
    □ 5
    □ 6
    □ ….greater than [last number] **Questions 2-9 apply only to your most recent working group.**
2. Please identify the funding agency or Synthesis Center for your most recent working group ():

> Pulldown box =
>
> List of 13 synthesis centres
>
> Other funding source
>
> Unfunded
>
> If “other funding source”, please specify: [text box]
3. What was your academic status when you participated most recently in a working group? [graduate student; postdoc; assistant professor, associate professor; professor; retired professor; non-academic researcher; other] If other, please specify [textbox].
4. How did you become involved in this working group? Textbox: I organized the group. I was invited to participate I applied to participate [if invited, path to 5:]
5. If you were invited to participate, how did you know the person who invited you? He/she was my graduate or postdoctoral supervisor (an organizer/participant) He/she was a previous collaborator. I knew the person although he/she was not a previous collaborator. I did not previously know the person who invited me. If other, please specify [textbox]
6. How many participants were in this working group? Of these participants, how many did you know prior to the working group?
7. There are many things that people do as part of working groups. Please check all activities that you have done as part of these groups. (Check all that apply).
  □ Compiled previously collected data
  □ Statistically analyse data
  □ Develop a mathematical model
  □ Run simulations
  □ Develop verbal model or framework
  □ Primary author of journal articles
  □ Edit journal articles
  □ Led a team of researchers
  □ Engaged in media work
8. How many participants from this working group have you subsequently collaborated with in other contexts?
9. Did this working group require the construction of a dataset? [if yes, go to:] Has this dataset been used for other projects, besides the original project of the working group?
10. Has your participation in this working group led to funding opportunities?
11. Have you ever declined an invitation to participate in a working group? Yes or No If yes, why: I was too busy or could not travel due to work-related duties I was too busy or could not travel due to family-related duties I was not interested in the topic I do not like the working group method Other If other, please specify [textbox] Thank you for participating in this research. If you would like more information on this study or to have a summary of our findings, please leave your email address. [textbox] We are offering a free CIEE mug to the first 100 researchers to complete this survey, which can be picked up by yourself (or someone you nominate) at the CSEE meeting in Fredericton Aug 18-21, 2019. If you would like to be considered for this, please leave your email here: [textbox]

